# Mmp14 is required for matrisome homeostasis and circadian rhythm in fibroblasts

**DOI:** 10.1101/2022.04.01.486699

**Authors:** Ching-Yan Chloé Yeung, Richa Garva, Adam Pickard, Yinhui Lu, Venkatesh Mallikarjun, Joe Swift, Susan H. Taylor, Jyoti Rai, David R. Eyre, Mayank Chaturvedi, Yoshifumi Itoh, Qing-Jun Meng, Cornelia Mauch, Paola Zigrino, Karl E. Kadler

## Abstract

The circadian clock in tendon regulates the daily rhythmic synthesis of collagen-I and the appearance and disappearance of small-diameter collagen fibrils in the extracellular matrix. How the fibrils are assembled and removed is not fully understood. Here, we first showed that the collagenase, membrane type I-matrix metalloproteinase (MT1-MMP, encoded by *Mmp14*), is regulated by the circadian clock in postnatal mouse tendon. Next, we generated tamoxifen-induced *Col1a2-Cre-ERT2::Mmp14* KO mice (*Mmp14* conditional knockout (CKO)). The CKO mice developed hind limb dorsiflexion and thickened tendons, which accumulated narrow-diameter collagen fibrils causing ultrastructural disorganization. Mass spectrometry of control tendons identified 1195 proteins of which 212 showed time-dependent abundance. In *Mmp14* CKO mice 19 proteins had reversed temporal abundance and 176 proteins lost time dependency. Among these, the collagen crosslinking enzymes lysyl oxidase-like 1 (LOXL1) and lysyl hydroxylase 1 (LH1; encoded by *Plod2*) were elevated and had lost time-dependent regulation. High-pressure chromatography confirmed elevated levels of hydroxylysine aldehyde (pyridinoline) crosslinking of collagen in CKO tendons. As a result, collagen-I was refractory to extraction. We also showed that CRISPR-Cas9 deletion of *Mmp14* from cultured fibroblasts resulted in loss of circadian clock rhythmicity of period 2 (PER2), and recombinant MT1-MMP was highly effective at cleaving soluble collagen-I but less effective at cleaving collagen pre-assembled into fibrils. In conclusion, our study shows that circadian clock-regulated *Mmp14* controls the rhythmic synthesis of small diameter collagen fibrils, regulates collagen crosslinking, and its absence disrupts the circadian clock and matrisome in tendon fibroblasts.

## INTRODUCTION

Tendon development in mice is apparent from embryonic day 14.5 when narrow (∼ 50 nm diameter) collagen fibrils begin to appear in channels between embryonic fibroblasts ^1^. The fibrils increase in number during embryonic development and remain attached to fibripositors at the cell surface. At birth, fibripositors disappear, and the fibrils detach from cells and increase in diameter ^2^. Thus, at the ultrastructural level, tendon development can be defined by stage 1 (embryonic) characterized by the presence of exclusively narrow fibrils, progressing to stage 2 shortly after birth when the fibril diameter distribution is multimodal (mean diameters of ∼ 50 nm, ∼150 nm and ∼250 nm). Mice lacking *Mmp14* (encodes MT1-MMP) are morphologically normal at birth but later exhibit reduced size and several abnormalities, and die within a few weeks of birth^3, 4^. Electron microscopy shows that detachment of the narrow fibrils from the cells and subsequent increase in fibril diameter do not occur in *Mmp14*-deficient neonatal mice ^2^; thus, tendon development arrests at the end of stage 1. A motivation for the present study was to determine the role of MT1-MMP in adult tendon ECM homeostasis. Of particular interest to the present study, the narrow fibrils in postnatal tendon are under the control of the circadian clock ^5^.

MT1-MMP is a transmembrane matrix metalloproteinase first identified for its ability to activate progelatinase A (proMMP2) and thereby enhance cellular invasion of reconstituted basement membrane ^6^. Extensive studies have showed that it can cleave a wide range of cellular and extracellular substrates including protease inhibitors, chemokines, cytokines, cell receptors, proMMP13, fibronectin and collagens (see ^7^). Therefore, aberrant *Mmp14* expression may result in major disruptive effects on matrix homeostasis. In humans, *Mmp14* has been implicated in cancer cell invasion ^8, 9^, which has been attributed to its ability to degrade extracellular matrix (ECM) macromolecules, especially type I collagen (collagen-I) ^10-12^. Like other vertebrate collagenases, MT1-MMP cleaves members of the fibrillar collagen family at the characteristic ¾-¼ site between residues 775 and 776 in each collagen α-chain ^13^. MT1-MMP is distinct from other MMPs in that its deletion in mice results in early postnatal death ^3, 4^. The mice fail to thrive and exhibit a wide range of pathologies in addition to walking difficulties including defective osteogenesis and angiogenesis ^3, 4, 14^, altered white adipose tissue ^15^ and modulation of inflammatory responses ^16^, activation of transforming growth factor β1 (TGFb1) ^17^, and deregulation of α5β1 integrin endocytosis ^18^.

We noted that the organ systems affected in the *Mmp14*-null mouse overlap with those affected in mice that lack a functional circadian clock. These include, bone formation ^19^, adipogenesis ^20^, inflammatory immune responses ^21^, and angiogenesis ^22^. Furthermore, as noted above, the narrow fibrils in postnatal mouse tendon are under circadian clock control. Circadian clocks are cell-autonomous time keeping mechanisms that optimize cellular activities in anticipation of varying time-of-day demands on the cell ^23, 24^. The molecular pacemaker is a transcription-translation feedback loop that is driven by two transcription activators, BMAL1 (bone and muscle Arnt-like 1) and CLOCK (circadian locomotor output cycles kaput protein), and two transcription repressors, CRY (cryptochrome) and PER (period) that inhibit the transcriptional complex^25^. A search of public databases showed that *Mmp14* transcripts are under circadian clock control in mouse suprachiasmatic nucleus (the central pacemaker located in the anterior hypothalamus), lung, heart, cartilage, and adipose tissue ^26^. Time-series transcriptomic profiling of mouse tendon (ArrayExpress accession no. E-MTAB-7743 ^27^) showed circadian expression of the *Mmp14* transcript. However, the biological role of a circadian-regulated matrix metalloproteinase in adult tendons is not fully understood.

Here we generated a type I collagen-specific conditional *Mmp14* knockout (CKO) mouse (*Col1a2*^*CreERT2*^::*Mmp14*^F/F^) and performed electron microscopy, proteomics, collagen content, and crosslink analyses of tendon. Our results show a circadian control of MT1-MMP in tendon and identify new functions for Mmp14 in maintaining a functional circadian clock in fibroblasts, in matrix homeostasis, and in ensuring normal crosslinking of collagen fibrils, in tendon.

## RESULTS

### *Levels of Mmp14* and MT1-MMP are circadian in postnatal mouse tendon

The readouts from 3 independent microarray probe sets showed the highest expression of *Mmp14* at the night-day transition (**Fig. S1A**, data replotted from data published by Yeung et al. 2014 ^27^). Time-series western blot analysis showed peak levels of MT1-MMP at circadian time 3 (CT3; 3 h into the day), which was approximately 4 hours after the peak in transcript levels (**Fig. S1B**). Next, we showed that time-of-day-dependent expression of Mmp14 observed in mouse Achilles and tail tendons was abolished in tendons from tendon-specific *Bmal1* knockout *(Scx-Cre::Bmal1*^*F/F*^) and global Clock-null (*Clock*Δ19) mice which lack circadian rhythms in their tendons ^5, 27^. We did this by examining relative *Mmp14* mRNA levels in tendons at two time points based on the peak and nadir *Mmp14* expression and MT1-MMP levels in mouse tendon: zeitgeber time 3 (ZT3; 3 h into an external or environmental cue, and in this case it was light of a 12-h light/12-h dark cycle) and ZT15 (3 h into the dark phase) (**Fig. S1C-E**). As such, the rhythmic expression of *Mmp14* depends on an intact local circadian clock in tendon tissues.

### Accumulation of narrow-diameter collagen fibrils in tendons with a postnatal deletion of *Mmp14*

Because global knockout of *Mmp14* results in perinatal death and the tendon-specific *Mmp14* knockout mouse (*Scx-Cre::Mmp14*^*F/F*^) does not progress past stage 1 tendon development ^2^, we generated the conditional collagen-I-specific *Mmp14* knockout mouse *(Col1a2*^*CreERT2*^*::Mmp14*^*F/F*^, abbreviated to *Mmp14* CKO or CKO), which was inducible by tamoxifen ^28^. We treated 4 week-old *Col1a2*^*CreERT2*^*::Mmp14*^*F/F*^ (abbreviated to *Mmp14* CKO or CKO) mice (time point 0, T0) with tamoxifen for 5 weeks to induce deletion of *Mmp14* in all cells expressing *Col1a2* (i.e., mainly fibroblasts including tenocytes) and harvested tendons at 10-, 14-, 18- and 22-weeks of age (T1, T2, T3 and T4, respectively; see schematic in **Fig. 1A**). We noticed that tamoxifen-treated mice had difficulty walking, severe skin thickening^28^ and dorsiflexion of the hind limbs (**Fig. 1B**). Examination of the tails showed thickening of tendon bundles (**Fig. 1C**). Q-PCR analysis of *Mmp14* expression in tendons at ZT3 and ZT15 showed time-dependent expression of *Mmp14* in wildtype (WT) mice but was almost undetected in tamoxifen-treated mice (**Fig. 1D**). Western blot analysis established there was an absence of MT1-MMP in primary fibroblasts isolated from *Mmp14* CKO tendons (**Fig. 1E**). Transmission electron microscopy analysis showed a progressive increase in the abundance of narrow-diameter (∼ 50 nm) collagen fibrils during T1 to T4 in CKO tendons compared to controls (**Fig. 1F, G** and **Fig. S2**). In T4 CKO tendons narrow fibrils were also observed within fibricarriers (**Fig. S2B**). We also noted a gradual increase in irregular fibril cross sections from T0 to T4 in CKO tendons (**Fig. 1H**). Serial block face-scanning electron microscopy, using established methods ^29^, showed gross disruption of the collagen fibril network in CKO tendons compared to tendons of littermates not treated with tamoxifen (**Fig. 1I, J** and **Movie 1** and **Movie 2**). During dissection we noticed that CKO tendons were brittle and would break easily. Therefore, samples were not analyzed by mechanical testing. These data confirm that *Mmp14* is necessary for postnatal tendon tissue homeostasis.

**Figure 1.**
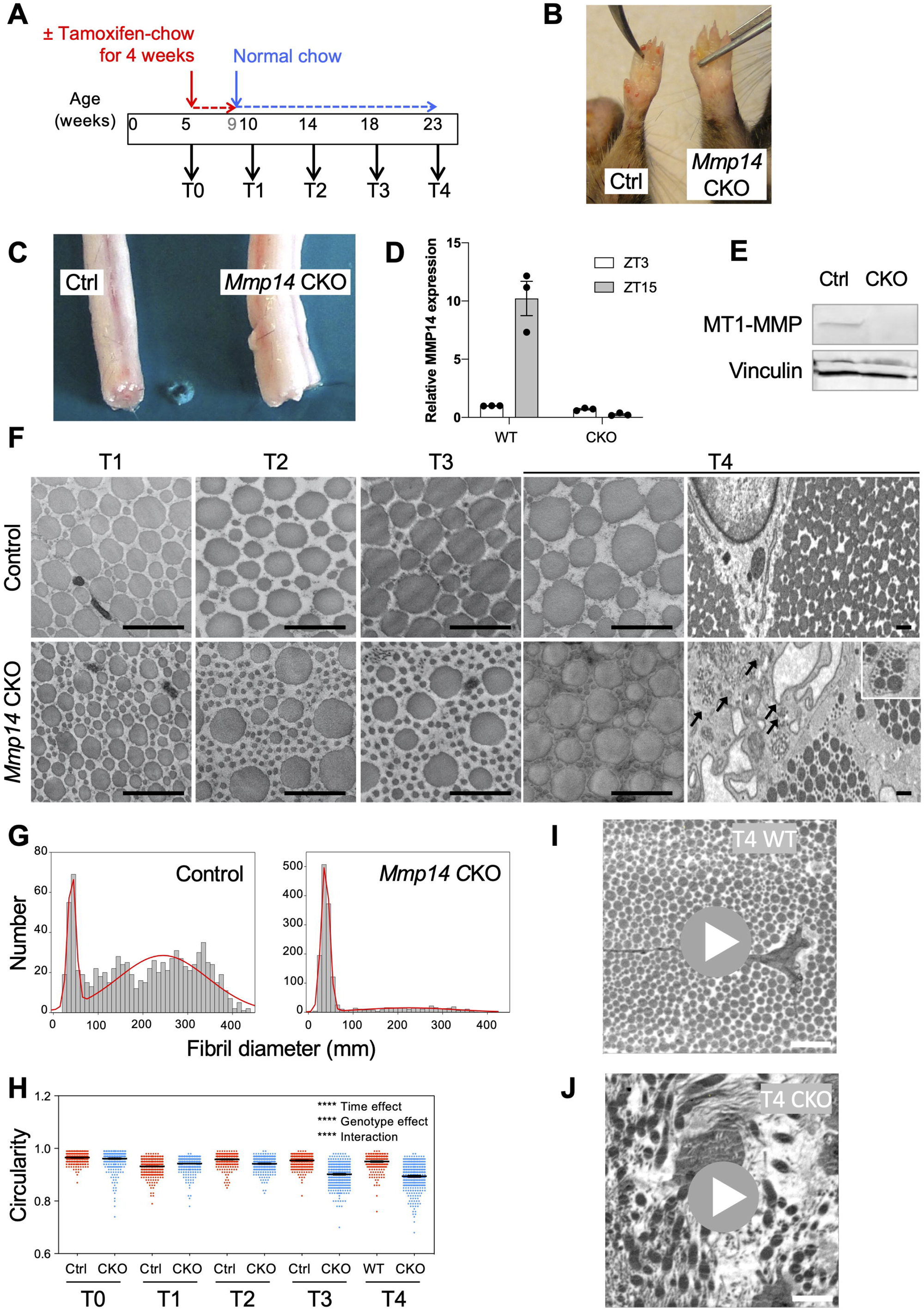
Accumulation of collagen fibrils in tendons from tamoxifen-induced conditional deletion of *Mmp14* of *Col1a2CreERT2:Mmp14*^*F/F*^ mice. **A,** Schematic of *Mmp14* deletion in fibroblasts of *Col1a2CreERT2:Mmp14^F/F^ (Mmp14 CKO)* mice in vivo by tamoxifen-chow feeding at 4 weeks of age for 5 weeks. Tissue was collected at T0 (before tamoxifen induction), T1, 2, 3 and 4 (1, 5, 9 and 13 weeks after tamoxifen induction, respectively). Control mice were not treated with tamoxifen. **B-C,** Photographs showing thickening of skin of *Mmp14* CKO at site-matched locations of the limb, and **C,** thickening of tendons in the tail compared to control litter mates at T4. **D,** Q-PCR analysis for *Mmp14* at T4 control and CKO tendons at ZT3 and ZT15 (n=3). **E,** Western blotting of MT1-MMP in fibroblasts isolated from control (Ctrl) and CKO Achilles tendons. Vinculin was used as a protein loading control. **F,** Transmission electron microscopy analysis showed *Mmp14* CKO tendons accumulated narrow diameter fibrils both in the ECM and inside cells (black arrows and inset in T4 panel) but not in control tendons. **G,** Fibril diameter distributions of fibrils measured in control and *Mmp14* CKO tendons at T4 show an increase in narrow-diameter (∼50 nm) fibrils in *Mmp14* CKO tendons. **H**, Circularity of collagen fibril cross sections of *Mmp14* CKO tendons showed significant irregularity compared to control fibrils at all time points after tamoxifen induction, with increasing irregularity over time (270-470 fibrils measured per time point; ****P<0.0001 for time, genotype and interaction). **I-J,** Single images selected from step-through Movie 1 and Movie 2 generated from volume-EM analysis of control and CKO Achilles tendons. Scale bar, 1 µm.

### Altered temporal patterns of abundance of soluble proteins in *Mmp14* CKO tendons

To understand the consequences of *Mmp14* deficiency on time dependent expression of proteins *in vivo* we performed proteomics analysis of tendon harvested at two time points in the 24-hour day from *Mmp14* CKO mice and control littermates (**Fig. 2A**). Note that, severe walking difficulties in *Col1a2-Cre-ERT2::Mmp14*^*F/F*^ mice treated with tamoxifen meant that insufficient numbers of mice were available for a circadian time series collection of tissues every 4 hours over two days as performed previously for studies of the tendon circadian clock ^5^. Instead, two time points, ZT3 and ZT15 were selected (**Fig. S1A** and **B**). T2 (5 weeks post tamoxifen treatment) was chosen because it was the time point where a more prominent fibrotic phenotype was detected in skin^28^ and tendons, deletion of *Mmp14* was complete, and it was still possible to dissect CKO tendons cleanly from the limb ^28^. We extracted non-covalently bound proteins with sodium laurate-sodium deoxycholate (SL-DOC) buffer using methods previously described ^5^. Analysis of samples identified 1195 proteins with ≥ 2 peptides per protein of which 212 showed time-dependent abundance, as shown by MetaCycle analysis, with good reproducibility between biological replicates as indicated by Pearson analysis across consecutive 24-hour periods (**Fig. S3A** and **Table S1 and Table S2**). Comparing logged ZT15/ZT3 fold changes for 1195 proteins showed that the absence of *Mmp14* in CKO tendons disrupted time-dependent protein abundances (**Fig. S3B**). Of the 212 time-dependent proteins detected in control tendons, 17 remained time-dependent and have a statistically significant (FDR adjusted p-value <0.05) difference in expression between the two time points in CKO tendons, 19 proteins exhibited a reversed time-dependent expression pattern (**Fig. 2B**), and 176 had lost time-dependent abundance pattern in CKO tissues (**Fig. 2C**). A search through the previously published rhythmic soluble mouse tail tendon proteome (see ^5^) revealed 27 common proteins amongst the 176 that had lost temporal abundance. Interaction analysis of these 27 proteins that have lost circadian rhythmic abundancy revealed direct interactions between fibronectin (FN1), the MT1-MMP substrate accumulated in *Mmp14*-deficient mice ^2^, and 19 out of 22 of the proteins recognized by Pathway Commons (**Fig. 2D**). Together, these data demonstrate that deletion of *Mmp14* in postnatal fibroblasts disrupts time-dependent abundance of soluble proteins, including those that have circadian rhythmicity, in tendon tissue.

**Figure 2.**
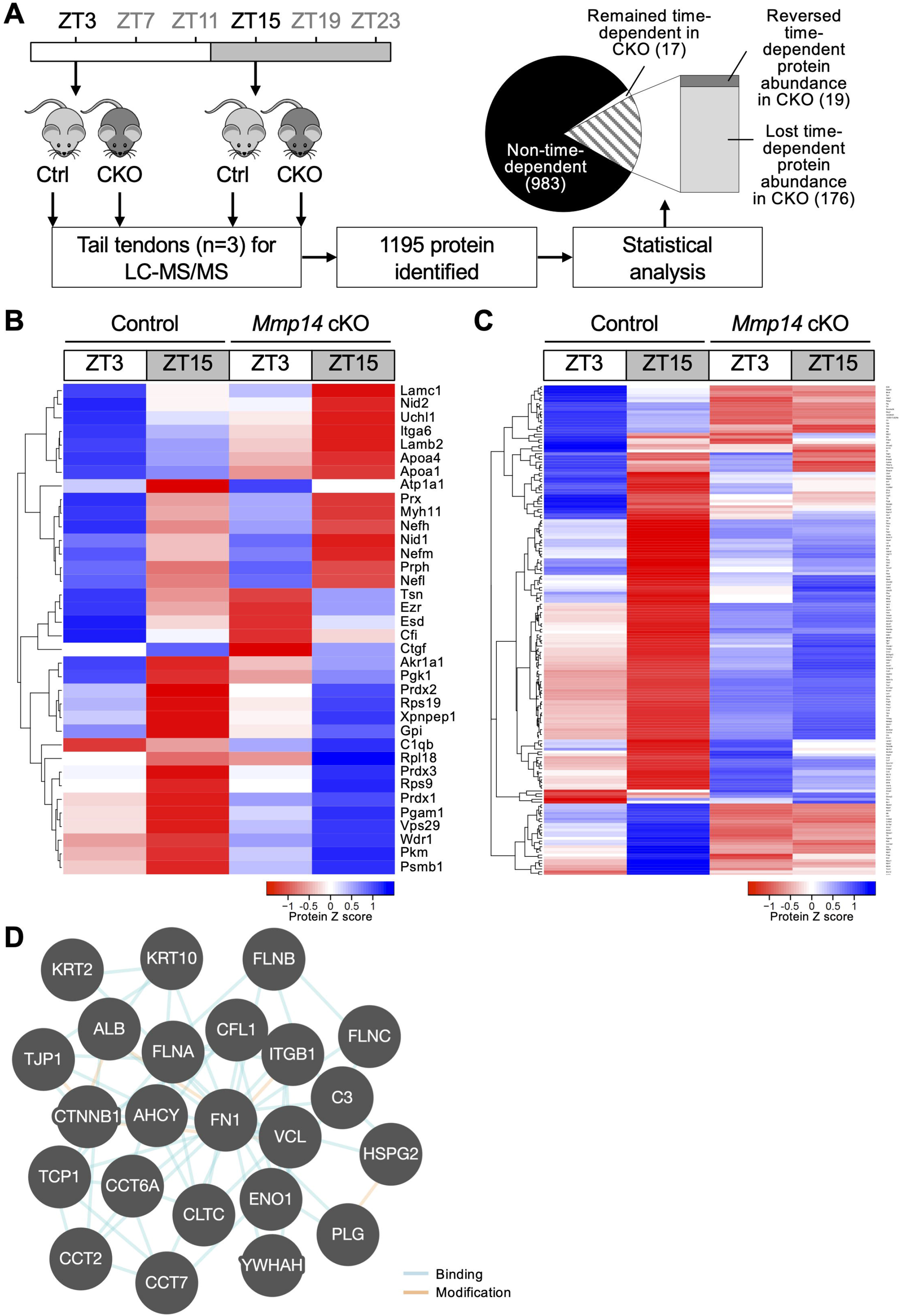
Proteomic analysis of tendons from conditionally-deleted *Mmp14* mice. **A**, Experimental strategy and results summary of proteomics analysis of protein abundance in tendons of *Col1a2CreERT2:Mmp14*^*F/F*^ (CKO) compared to control littermates at ZT3, when *Mmp14* levels are highest, and at 12 h apart (ZT15) (n=3). 1195 proteins were identified, 212 were identified as significantly time-dependent in control tendons (p<0.05), of which 17 remained time-dependent, 19 showed a statistical significant reversal of abundance pattern, and 176 were no longer time-dependent in *Mmp14* CKO tendons. **B**, Heat map representation of proteins that remained time-dependent in *Mmp14* CKO tendons, including proteins with abundance in anti-phase to wild type tendons. **C**, Heat map representation of the abundance of 176 proteins that are time-dependent in control tendons but are no longer time-dependent in *Mmp14* CKO tendons. **D**, Interaction analysis of proteins that have lost circadian rhythmic abundancy in CKO tendons.

### Dysregulation of the tendon matrisome and elevated crosslinking in the absence of *Mmp14*

Since *Mmp14* CKO tendons exhibit gross ultrastructural changes in the ECM an investigation of matrisome proteins was extended to proteins that did not have time dependent abundance. We found that 121 matrisome proteins had differential abundances in CKO tendons compared to control tendons (i.e., 62 proteins higher and 59 proteins lower abundance in CKO tendons; **Fig. 3**). Interactions analysis revealed the direct and indirect effects of *Mmp14* deletion on and via substrates and binding partners (**Fig. S4**). For example, there is increased abundance of the MT1-MMP14 substrate proMMP2, which may have caused the modulated abundances of fibronectin, cathepsin K (CTSK, which is circadian clock regulated and involved in collagen turnover), collagen V (COL5A1), collagen VI (COL6A1, COL6A2), collagen XIV (COL14A1), collectin-12 (COLEC12), galectin 3 (LGALS3), tenascin C (TNC), tissue inhibitor of metalloproteinase 2 (TIMP2), and thrombospondin 2 (THBS2). Not all proteins were increased in abundance, e.g., there were reductions in COL2A1, COL6A2, COL6A3, and decorin (DCN). Of particular interest, enzymes that catalyze posttranslational modifications in collagen leading to crosslink formation were elevated and included lysyl hydroxylase 2 (also known as procollagen-lysine, 2-oxoglutarate 5-dioxygenase 2 (PLOD2)) (up 13-fold), and lysyl oxidase-like 1 (LOXL1) (up 20-fold).

**Figure 3.**
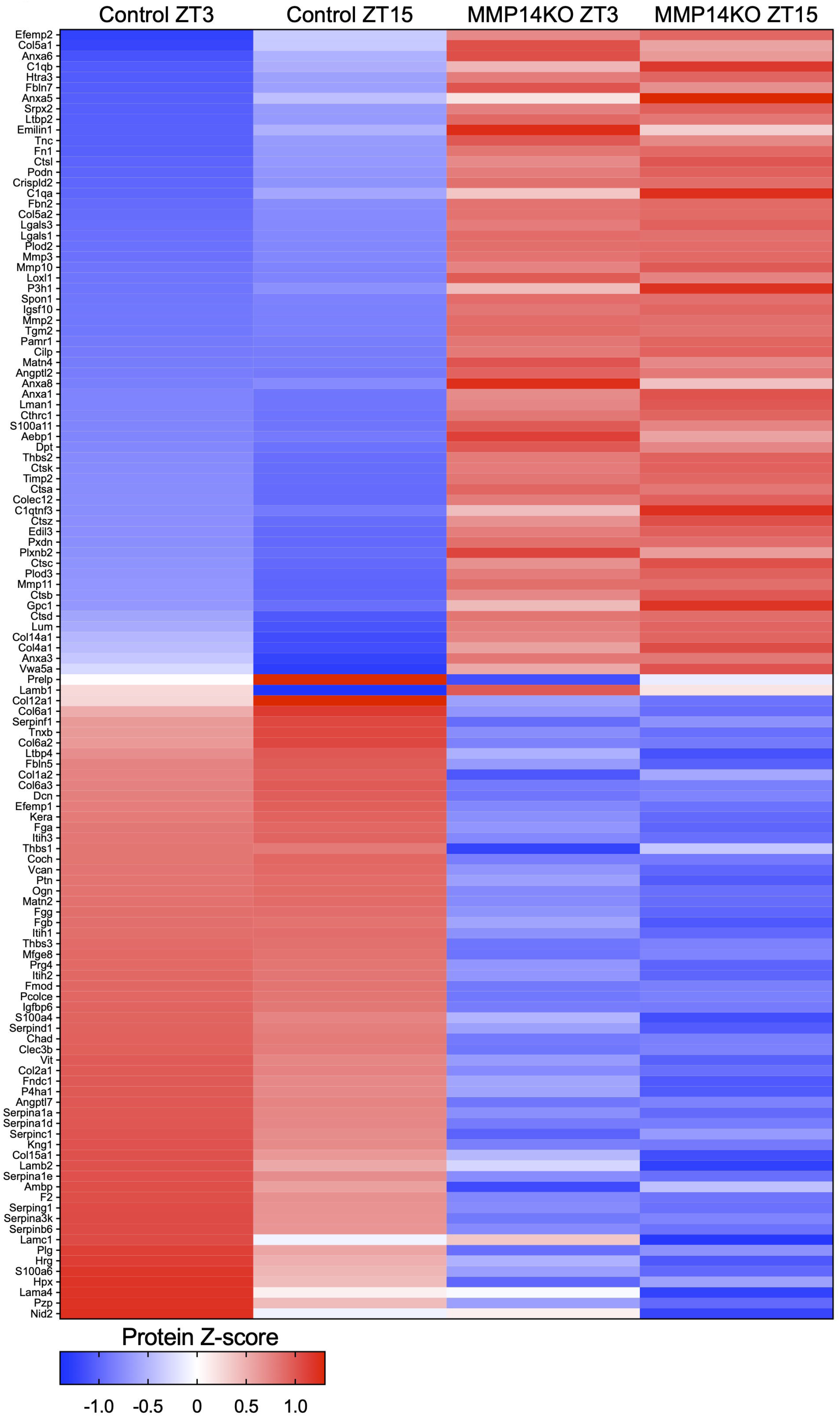
Disruption of the soluble matrisome in CKO tendons. Heatmap of Z-scored fold changes for matrix proteins with differential abundance between control and CKO tendons (62 increased abundance, 60 decreased abundance). Proteins are listed ranked by increasing abundance in control tendons at ZT3.

There was clear discrepancy between the accumulation of collagen fibrils in the *Mmp14* CKO mice (see **Fig. S2B**) and the low levels of collagen-I detected by mass spectrometry of CKO tissues (**Table S1** and **Fig. S4C**). Therefore, we collected tendons from T2 *Mmp14* CKO and control mice at ZT3 and ZT15 and performed hydroxyproline analysis to quantify collagen. We also subjected tendons to extraction in SL-DOC buffer, followed by hydroxyproline quantification. Analysis of whole tissue and SL-DOC extracts showed ∼98%:2% split between crosslinked collagen in whole tissue and soluble collagen in the SL-DOC extract (**Table 1**). There was an increase of ∼ 6.4% in overall collagen content in CKO tendons, which was largely the result of collagen accumulated at ZT15. However, there was ∼8-fold increase in the amount of hydroxylysine aldehyde in *Mmp14* CKO compared to control tendons at both time points (**Table 1**). Mass spectrometry ruled out significant amounts of collagen-III as the source of hydroxylysine aldehyde in the CKO samples. Therefore, the discrepancy in the number of collagen-I peptides extracted into SL-DOC buffer, and the marked accumulation of collagen fibrils in the CKO tendons, was the result of accumulation of collagen at ZT15 and decreased extractability because of increased hydroxylysine aldehyde cross-linking of collagen-I ^30^ in the CKO tissues.

**Table 1.** Collagen content in control and *Mmp14 C*KO tendons determined by hydroxyproline assay. N = 3. Values are mean ± SEM. Two-way ANOVA: *p=0.0146 for time effect, **p=0.0014 for genotype effect.

|  | Control |  | CKO |  |
| --- | --- | --- | --- | --- |
|  | ZT3 | ZT15 | ZT3 | ZT15 |
| % hydroxyproline content per lyophilized whole tissue weight | 70.97 ± 4.27 | 62.30 ± 5.25 | 71.43 ± 1.74 | 70.40 ± 6.35 |
| % hydroxyproline content of SL- | 1.50 ± 0.10 | 0.98 ± 0.31* | 1.39 ± 0.12 | 0.81 ± 0.05* |
| Hydroxylysine aldehyde (pyridinoline) content per whole tissue hydroxyproline | 0.009 ± 0.003 | 0.013 ± 0.002 | 0.095 ± 0.033** | 0.095 ± 0.007** |

### Elevated *Pitx2* and PITX2 nuclear localization in *Mmp14* knockout cells

To understand how loss of *Mmp14* led to extensive crosslinking, we investigated a possible relationship between *Mmp14* and paired-like homeodomain transcription factor 2 (PITX2), which is a transcription factor that has been reported to regulate the expression of *Plod2* ^31, 32^, as well as collagen I and V ^33^. We showed by Q-PCR that levels of *Pitx2* mRNA are low in murine tail tendon at both ZT3 and ZT15 but markedly elevated in the CKO tendons (**Fig. 4A**). *Plod2* expression was also elevated ∼10 fold in CKO tendons (**Fig. 4B**). Next, we expressed PITX2-GFP in immortalized tail tendon fibroblasts (iTTFs). Over-expression of PITX2-GFP was confirmed by Q-PCR (**Fig. 4C**), and although it was not as pronounced as in *Mmp14* CKO tendons, resulted in a 2-fold increase in expression of *Plod2* (**Fig. 4D**), confirming that PITX2 drives the expression of *Plod2* in tendon fibroblasts. PITX2-GFP was localized to cell nuclei (**Fig. 4E**) and this nuclear localization was enhanced in *Mmp14*-depeleted fibroblasts (**Fig. 4F**). Over-expression of mCherry-MT1-MMP in PITX2-GFP expressing iTTFs reversed the induced *Pitx2* and *Plod2* expression and enhanced *Col1a1* expression (**Fig. 4G-J**). Together, these data show that *Mmp14* negatively regulates *Plod2* expression via PITX2.

**Figure 4.**
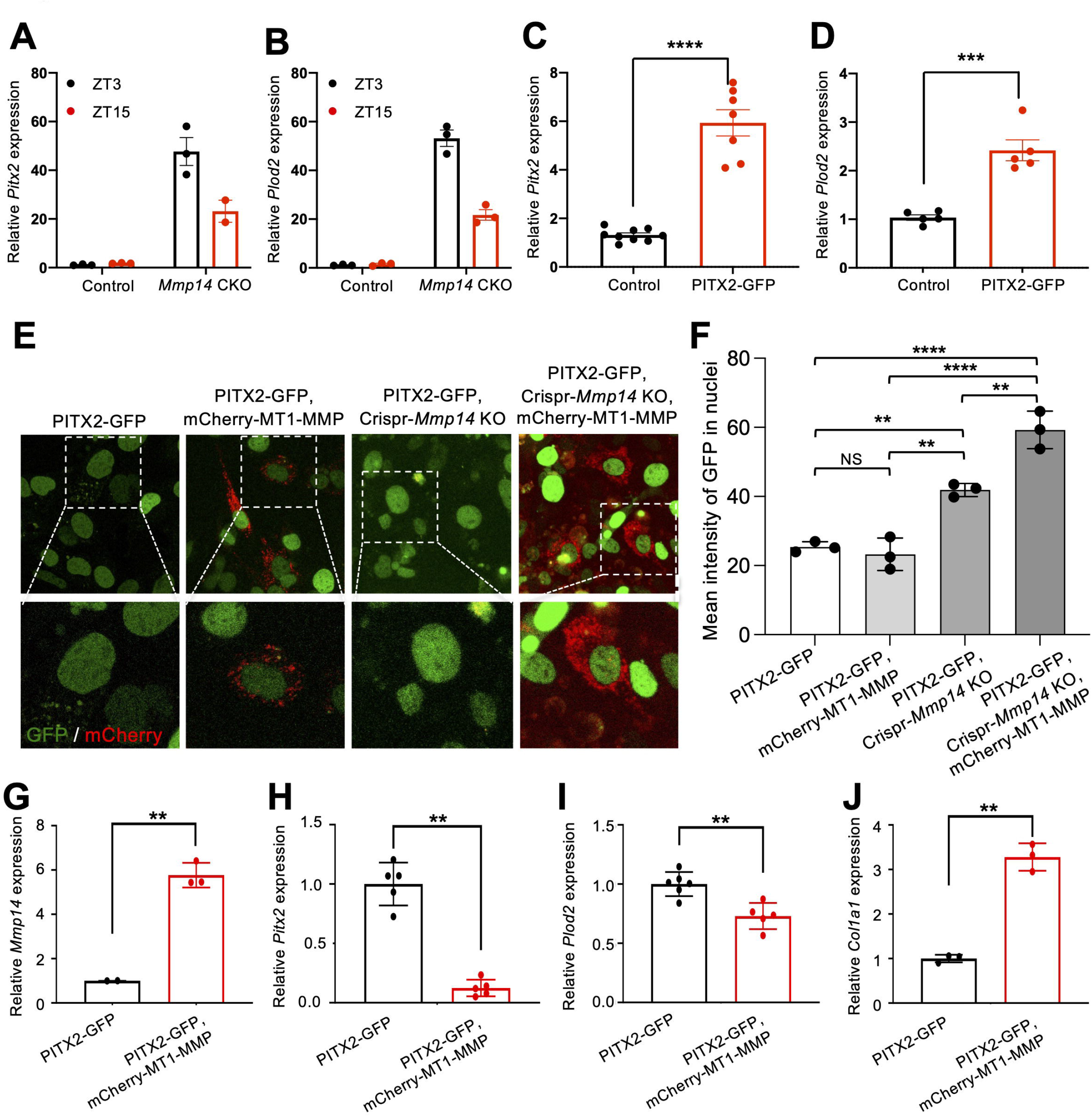
PITX2 Is elevated in *Mmp14* CKO tendons. **A**, Q-PCR detection of *Pitx2* and **B**, *Plod2* mRNA levels in control and *Mmp14* CKO tail tendons 5 weeks post tamoxifen treatment, at ZT3 and ZT15 (n=3). **C**, Per2::Luc iTTFs stably transduced with control or PITX22-GFP lentivirus (n=3), demonstrating enhanced expression of *Pitx2* (n=9 and n=7; ****p<0.0001) and **D**, a concomitant increase in *Plod2* mRNA expression (n=5; ***p<0.001). **E**, Live cell Airyscan imaging of Per2::Luc iTTFs and *Mmp14* KO ITTFs stably transduced with Pitx2-GFP lentivirus. Cells were transiently transduced with mCherry tagged MMP14 carrying a silent mutation in the CRISPR-Cas9 PAM site. Pitx2-GFP was readily detected within the nucleus but also with punctate localisation within the cell body in control cell, these correlated with regions also positive for MT1-MMP. Nuclear Pitx2-GFP signals were stronger within *Mmp14* KO cells and also displayed reduced localisation within the cell body. Re-expression of *Mmp14* within knockout cells reduced nuclear PITX2-GFP signals. **F**, Quantification of mean nuclear GFP signal intensity (n=3; ****p<0.0001, **p<0.01). **G-J**, Q-PCR demonstrating expression of *Mmp14* in PITX22-GFP expressing iTTFs, which resulted in reduced *Pitx2* and *Plod2* mRNA expression, but enhanced *Col1a1* mRNA (n=3; **p<0.01).

### MT1-MMP-mediated cleavage of collagen-I is dispensable for postnatal tendon development

The amassing of narrow collagen fibrils in the CKO tendons prompted us to investigate if their accumulation was a direct result of the absence of collagen fibril turnover in the absence of MT1-MMP. Thus, we released intact collagen fibrils from 6-week-old mouse Achilles tendon and examined the diameters before and after incubation of the fibrils with recombinant soluble MT1-MMP (rMT1-MMP), which was synthesized as previously described ^34^. The fibril diameters were derived from mass-per-length measurements using scanning transmission electron microscopy ^35^. The results showed no differences in fibril diameters between rMT1-MMP treated and untreated samples (**Fig. 5A**). When assembled into fibrils, many of the collagen molecules are buried in the core of the fibril and thereby inaccessible to MT1-MMP. Therefore, we incubated fibrils that had been released from tendon overnight at 37 °C (to keep the fibrils intact) and 30 °C (to partially depolymerize the fibrils), and then added rMT1-MMP to the suspension of fibrils. In control experiments we incubated MT1-MMP with soluble collagen-I (i.e., not assembled into fibrils) in the presence and absence of EDTA, which chelates Ca2+ ions and inactivates MT1-MMP (**Fig. 5B**). SDS-PAGE and mass spectrometry were used to detect cleavage products. Liquid chromatography with tandem mass spectrometry (LC-MS/MS) analysis of the cleavage products showed the presence of a semi-tryptic peptide from the ¾ α1(I) fragment corresponding to the expected ¾-¼ cleave site, which confirmed that rMT1-MMP had cleaved soluble collagen-I molecules at the expected Gly775 and Ile776 site (**Fig. 5C** and **D**). However, incubation of rMT1-MMP with the suspension of fibrils resulted in only a small proportion of the collagen being cleaved and only when the reaction was performed overnight at 30 °C (**Fig. 5B**, white box). The most likely explanation was partial depolymerization of the fibrils at this temperature, resulting in release of collagen molecules that were then susceptible to cleavage by rMT1-MMP. As expected, cleavage was inhibited by EDTA. We noticed a downward shift in mobility of β-chains (crosslinked α-chains), which was not inhibited by EDTA. It is likely, therefore, that this was the result of non-MMP activity in the collagen fibril preparation (**Fig. 5B**). These results showed that collagen is largely protected from cleavage by MT1-MMP when it is assembled into fibrils.

**Figure 5.**
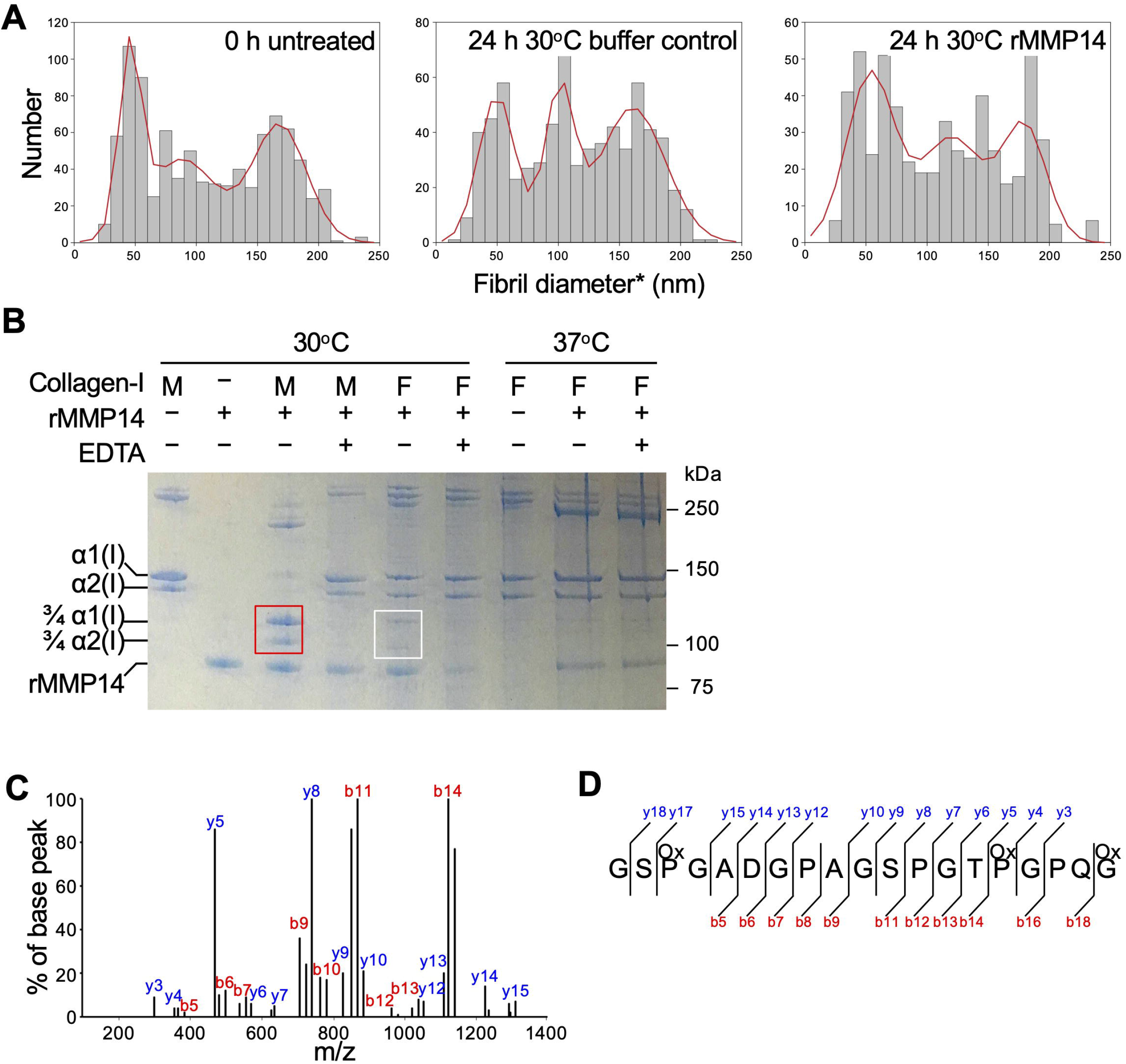
Recombinant MT1-MMP preferentially cleaves soluble collagen-I. **A**, Fibril diameter distribution of collagen fibrils dispersed from mouse Achilles tendons before treatment and 24 h after treatment with and without rhMT1-MMP at 30 °C. *Effective hydrated fibril diameter (>500 fibrils measured). **B** Coomassie blue-stained gel showing incubation of monomeric and fibrillar collagen-I with recombinant human MT1-MMP at 30 °C and 37 °C, in the presence or absence of 50 mM EDTA. M, monomeric; F, fibrillar collagen dispersed from Achilles tendon. rhMT1-MMP generated ¾α1(I) and ¾α2(I) fragments from monomeric collagen I (red box) and from fibrillar collagen (white box). Asterisk shows the b-crosslinked chains of collagen-I. A prominent faster migrating b-dimer is seen in the MT1-MMP treated samples which occurs in the presence and absence of 50 mM EDTA. **C**, Mass spectrum of rhMT1-MMP generated fragments from monomeric collagen I shows the mass to charge ratio of individual ions resulting from MS/MS fragmentation pattern between Gly_775_ and Ile_776_. **D**, Precise fragmentation pattern that contributes to the generation of b and y ions.

To investigate further the contribution of the collagen-I ¾-¼ cleavage site to collagen fibril homeostasis, we used transmission electron microscopy to examine tendons from the *Col-r/r* mouse, which carries a targeted mutation in *Col1a1* ^36^ that renders both the α1(I) and α2(I) chains resistant to cleavage by MMPs at the ¾-¼ site ^37^ (**Fig. S5A**). The results show that 23-week-old (equivalent to T4 *Mmp14* CKO mice) *Col-r/r* mouse tendons exhibit a broad distribution of collagen fibril diameters, in contrast to what was observed in the CKO mice (compare **Fig. 5B-E**). These observations showed that the accumulation of narrow fibrils in the postnatal CKO mice is not fully explained by the lack of cleavage of the ¾-¼ site in collagen-I.

### Deletion of *Mmp14* dampens the PER2::Luc circadian rhythm amplitude in cultured tendon fibroblasts

The occurrence of increased numbers of narrow fibrils and increased irregular fibril profiles were reminiscent of tendons with a disrupted circadian rhythm ^5^. To determine if *Mmp14* deletion affects the endogenous circadian clock, we used iTTFs from the circadian reporter PER2::Luc mouse (iTTF-PER2::Luc or abbreviated to ‘iTTFs’), which enabled us to monitor the circadian rhythm in real time ^5^. In initial experiments we incubated clock synchronized iTTF cells with the broad spectrum MMP inhibitor GM6001, which markedly reduced the amplitude of the circadian rhythm in treated cells compared to untreated controls (**Fig. S6A, B**). Next, we performed siRNA knockdown of *Mmp14*. Q-PCR showed that knockdown was highly effective at reducing the expression of *Mmp14* (**Fig. S6C**) and bioluminescence showed that the circadian clock rhythm was dampened in si*Mmp14* treated cells (**Fig. S6D**). Next, we performed clustered regularly interspaced short palindromic repeats (CRISPR) and CRISPR-associated protein 9 (Cas9) mediated knockout of *Mmp14* in iTTFs (**Fig. S6E**). Western blot analysis using an anti-MT1-MMP antibody showed absence of MT1-MMP in 7 different clones (**Fig. S6F**). *Mmp14*-null iTTF clones were unable to sustain a circadian rhythm in response to dexamethasone (**Fig. 6A**). Restoring *Mmp14* expression alone via lentiviral transduction rescued robust circadian oscillations (**Fig. 6B, C**). Together these data show that in mouse tendon tissue *Mmp14* expression is required for a robust circadian rhythm in tendon fibroblasts.

**Figure 6.**
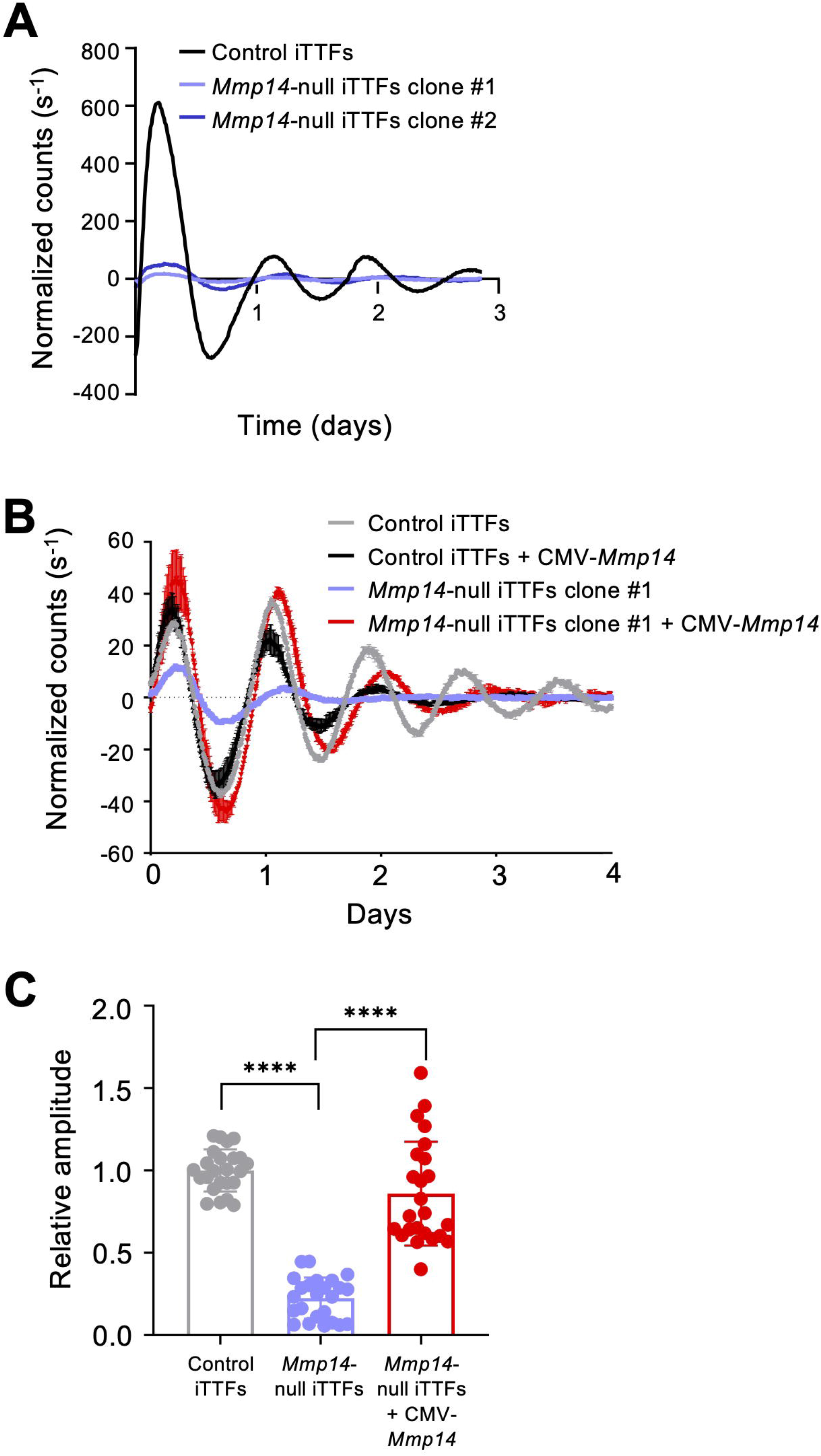
Deletion of Mmp14 weakens circadian entrainment. **A**, Representative bioluminescence recordings from Per2::Luc iTTFs (black) and single cell clones of CRISPR-Cas9 induced knockout of *Mmp14* Per2::Luc fibroblasts (blue) after synchronization with dexamethasone. **B**, Bioluminescence recordings from Per2::Luc iTTFs (grey) or *Mmp14* KO iTTFs (blue) that have been transduced with a lentivirus expressing *Mmp14* (carrying a silent mutation in the CRISPR-Cas9 PAM site) from a CMV promoter (black and red). Restoration of Mmp14 levels in *Mmp14* KO fibroblasts reinstated dexamethasone induced circadian rhythms. **C**, Relative amplitude of control iTTF cells, Clone #1 *Mmp14*-null iTTFs and Clone #1 *Mmp14*-null cells expressing *Mmp14* from a CMV promoter (n=4).

## Discussion

This study has shown that *Mmp14* is a master mediator of ECM homeostasis in adult tendons, as shown by the accumulation of small-diameter collagen fibrils and major changes in the matrisome when MT1-MMP is absent. We suggest that accumulation of the narrow collagen fibrils is the result of the accumulation of MT1-MMP substrates as well as loss of robust circadian rhythm, widespread changes to the ECM proteome, and elevated collagen crosslinking. Based on these findings we propose that a bi-directional dependency exists between matrix homeostasis and the circadian clock.

The lead observation for this study was the circadian expression of *Mmp14*. The peak of *Mmp14* expression in murine tendon is at CT3, which is in good agreement with the peak of *Bmal1* expression ^27^. Circadian clock regulated genes usually contain one or more transcriptional elements, RORE, DBPE and E-box ^38^. Noteworthy, a CACGTT (E-box) sequence is located 1.6 kb upstream of the translational start site of murine and human *Mmp14*, which would facilitate rhythmic binding of CLOCK-BMAL1 to E-box motifs ^39^. *Mmp14* gene expression has been shown to be rhythmically expressed in murine lung, heart, kidney and adipose as well as the suprachiasmatic nucleus ^40^. Therefore, it is likely that *Mmp14* is a BMAL1 responsive gene and its expression is restricted to a specific time of day across organ systems. We confirmed at the protein level that MT1-MMP is under circadian clock control in tendon, which was in line with circadian rhythmic levels of MT1-MMP and MMP2 activation observed in murine tendon fibroblasts ^41^. The peak of abundance of MT1-MMP was particularly narrow (∼ 4 hours after the peak of *Mmp14*) (**Fig. 1**), which is consistent with observations that MT1-MMP has a fast turnover of 4.8 h, and puts MT1-MMP in the top 1% of rapidly turned-over proteins in the proteome ^42, 43^. Together, these results and those of others studying organs other than tendon, show that *Mmp14* is under the control of the circadian clock which results in the synthesis of MT1-MMP in the morning. The appearance of MT1-MMP at CT3-7 coincides with the rhythmic abundance of collagen-I and with the onset of collagen fibril formation, in tendon ^5^, suggesting that *Mmp14* is important for collagen fibril formation.

Previously, a *Scx-Cre* driver was used to delete *Mmp14* in embryonic tendons ^2^ because *scleraxis* is expressed by all tendon progenitors ^44^. *Scx-Cre::Mmp14*^*F/F*^ neonatal tendons were thinner because the potential fibronectin ‘molecular bridge’ between the cell surface and collagen-I fibrils cannot be cleaved by MT1-MMP. So there is an overall reduction in fibrils assembled in the ECM, perhaps limited to cell size and/or fibril nucleation sites on the cell surface ^45^, contributing to the smaller tendon width ^2^. The use of an inducible *Scx-Cre* in postnatal tendons would not yield a complete *Mmp14* deletion because scleraxis expression in tendon becomes limited to the epitenon by 3 months of age ^46^. The major cell type in postnatal mouse tendons is the fibroblast and single cell transcriptomics experiments performed on healthy adult mouse Achilles tendons have confirmed that *Col1a2* expression is limited to fibroblast ^47^. Therefore, a tamoxifen-inducible *Col1a2-Cre* was chosen to drive the deletion of *Mmp14* in postnatal tendons. In contrast to thinner neonatal tendons resulting from embryonic *Mmp14* deletion, postnatal *Mmp14* deletion is initiated at 5 weeks of age, well into stage 2 of tendon development when collagen-I fibrils are released from the cell surface and permitted to grow laterally ^45^ and when a circadian rhythm is established in peripheral tissues in rodents (reviewed by ^48^). The consequence is tendon thickening, resulting from multiple factors: (1) halted turnover of fibronectin and collagen V, which act as nucleators for collagen-I fibril assembly, (2) lack of removal of circadian-regulated collagen-I that add onto existing large fibrils, (3) increased abundance of collagen crosslink enzymes (via PITX2), which stabilize new narrow fibrils and collagen-I molecules on larger fibrils, and (4) major disruption to the circadian rhythm and the temporally regulated matrisome, including molecules known to regulate collagen fibrillogenesis (e.g., lumican, etc).

We showed that tamoxifen-induced deletion of *Mmp14* in postnatal mice resulted in a severe tendon phenotype marked by dorsiflexion of hind limbs, tissue thickening, mistimed expression of 195 proteins (including 18 matrisome proteins) and altered abundance of a 104 matrisome proteins. The relatively large number of proteins with altered abundance is in line with ChIP and reporter vector studies showing that MT1-MMP regulates more than 100 genes ^49^. Our proteomics analysis of time-dependent abundance of proteins showed that MT1-MMP is a major regulator of the matrisome in tendon. Proteins with elevated levels included fibronectin, collagens I and V (which are major protagonists in fibril assembly in tendon), collagen XIV (its expression in tendon is associated with collagen fibril assembly and growth ^50, 51^), and prolyl 3-hydroxylase, whose absence in mice leads to tendon abnormalities ^52^. Fibronectin is a substrate for MT1-MMP; therefore, its increase in abundance might be a direct result of the absence of *Mmp14*. However, MMP3 (stromelysin-1), which can degrade fibronectin ^53^, was upregulated in CKO tendons. The abundance of fibronectin might also be in response to injury or inflammatory signals caused by the lack of *Mmp14*, which has been shown to put a brake on pro-inflammatory responses (reviewed by ^49^), perhaps by activating PI3/Akt3/GSK3 signaling ^16^. Notably, lumican was upregulated whereas the other small leucine rich proteoglycans, decorin, fibromodulin, keratocan, and lubricin, were down regulated. Lumican is a collagen-binding proteoglycan ^54^ and was proposed to facilitate interchain collagen crosslinking by interacting with collagen and lysyl oxidase (reviewed by ^55^). Lumican has a critical role in the organization of collagen fibrils in tendons and cornea and is profibrotic in heart ^56^. Furthermore, lumican knockout leads to reduced collagen crosslinking ^57^. The increased levels of lumican were matched by the increase in levels of LOXL1 and LH2, which post-translationally modify lysyl residues in collagen and lead to highly stable crosslinks, especially in collagen-I. Taken together, these protein changes are in line with increased pyridinoline crosslinks, which explains the low extractability of collagen-I from the CKO tendons.

We suggest that the circadian clock in *Mmp14*-null cells is ‘stuck’ at ZT15. Therefore, proteins that would normally be expressed only at night are expressed day and night in CKO tendons. This would explain the overabundance of pro-matrix proteins such as collagen-I, collagen-V, collagen-XIV, lumican, fibronectin, LOXL1, and LH2, which are normally only expressed at night but in the CKO tendons are expressed throughout 24 hours (see **Fig. 3**). The reasons why the small diameter fibrils are attached to cell surfaces, perhaps via fibripositors, is unclear. One explanation is that the processes of fibril turnover and internalization are arrested in CKO tendons. The observation that collagen assembled into fibrils is mostly inaccessible to MT1-MMP might suggest that fibril turnover is not the primary explanation. Alternatively, the accumulation of fibrils might result from the inability of cells to release collagen fibrils from the cell surface. As previously suggested, fibronectin might act as a molecular bridge between collagen and receptors at the cell surface ^2^. The appearance of a more normal collagen matrix in the *Col-r/r* mouse would also suggest that lack of cleavage of collagen-I (at least at the ¾-¼ site) by MT1-MMP does not explain the accumulation of narrow fibrils in CKO tendons. The persistence of collagen fibrils in the absence of MT1-MMP may be partially explained by an increase in covalent crosslinks in collagen (via LOXL1 and LH2, and promoted by lumican). Moreover, the increased availability of soluble collagen-I, in the absence of MT1-MMP, would be expected to lead to an increase in the number of collagen fibrils formed. Taken together, we propose that the accumulation of small diameter collagen fibrils attached to surfaces of tendon resident cells in CKO tendons is likely to be the consequence of compounding factors including the circadian clock being stuck at ZT15, wholesale changes in the matrisome, lack of proteolytic cleavage of collagens, fibronectin and other matrisome proteins, increased collagen crosslinking (making the fibrils refractory to disassociation and turnover), increased synthesis of collagen-I and V, and failure of cells to release fibrils to the matrix. In conclusion, we have demonstrated the pivotal and multifaceted role of *Mmp14* in circadian clock biology, matrisome regulation, and collagen fibril formation, together with the maturation of tendon fibroblasts during early postnatal growth that requires *Mmp14*. Our data also highlight the interdependence of Mmp14-dependent matrix homeostasis and the circadian clock.

Finally, our proteomics analysis identified additional changes in the proteome not directly related to the matrisome (data shown in **Fig. 2C** and **Supplementary Data 2**). These included changes to the cytoskeleton, which our current study could not distinguish between direct or indirect consequences of the absence of *Mmp14*. It has been reported that the cytoplasmic and transmembrane domains of MT1-MMP regulate non-proteolytic functions, including signal transduction via the cytoplasmic tail activating ERK1/2 signaling independent of proteolysis ^58, 59^, and gene regulation via a presumed nuclear localization signal in the transmembrane domain (reviewed by ^60^). MT1-MMP has a presumed nuclear localization signal ^49, 61^, and localizes to other intracellular sites including caveolae, mitochondria, and the cytoskeleton to regulate calcium levels leading to contractile dysfunction, and altered gene expression and ribosomal RNA transcription (reviewed by ^60^). It is possible that MT1-MMP has an intracellular scaffolding function that regulates collagen fibril formation and matrix homeostasis.

## Acknowledgements

The research was funded by Wellcome (110126/Z/15/Z and 203128/Z/16/Z) and the BBSRC (BB/T001984/1) to KEK. CM and PZ were supported by Deutsche Forschungsgemeinschaft through SFB 829 (Project-ID 73111208-SFB829 (B4)). VM was supported by a studentship from the Sir Richard Stapley Educational Trust. JS was funded by a Biotechnology and Biological Sciences Research Council (BBSRC) David Phillips Fellowship (BB/L024551/1). BC is supported by a Wellcome PhD studentship (210062/Z/17/Z). MC and YI was funded by Polish Ministry of Science & Higher Education (1291/MOB/IV/2015/0). Q-JM was supported by Medical Research Council project grants MR/T016744/1 and MR/P010709/1 and Versus Arthritis Senior Research Fellowship Award 20875. National Institutes of Health (NIH) Grant: (NIAMS) AR037318 to DRE. The authors would like to thank Ronen Schweitzer (Shriners Hospital for Children, Portland, OR, USA) for Scx-Cre mice, Jan Zamek, Alison Hallworth and Raymond Hodgkiss, and the University of Manchester Biological Support Facility for assistance in animal welfare and husbandry. Re-derivation of mouse strains and generation of expression vectors were performed by the Genome Editing Unit, University of Manchester. The proteomics was performed at the Biological Mass Spectrometry Facility in the Faculty of Biology, Medicine and Health (University of Manchester) with the assistance of Stacey Warwood and Ronan O’Cualain, and electron microscopy was performed in the Electron Microscopy Facility, Faculty of Biology, Medicine and Health (University of Manchester).

## Author contributions

Conceived the project and provided mechanistic insight: AP, C-YCY, KEK, RG.

Interpretation of results regarding the circadian clock: Q-JM.

Finalized the manuscript: C-YCY, KEK.

Designed and performed experiments and interpreted data: AP, C-YCY, DE, JR, MC, PZ, RG, ST.

Provided reagents: CM, PZ, YI.

Performed the electron microscopy: YL.

Performed bioinformatics analysis: JS, VM.

## Supplementary information

**Figure S1. *Mmp14* expression and MT1-MMP translation in tendon are under circadian clock control. A**, Expression of *Mmp14* mRNA in WT mouse tail tendons analyzed by microarray every 4 h during 48 h, relative to the first time point. Three colored lines indicate the results from 3 probe sets on the microarray chip, n=2; CT, circadian time (free-running time, in constant darkness). **B**, Levels of MT1-MMP in WT mouse tail tendons analyzed by western blotting. Levels of GAPDH protein served as a loading control. **C**, Relative expression of *Mmp14*, measured at zeitgeber time 3 (ZT3; 3 hours into the light phase) and ZT15 in mouse Achilles and tail tendons of WT mice (n=3 and **p=0.0045 for Achilles tendon; n=4 and **p=0.0093 for tail tendon), and **D**, tendon-specific Bmal1 knockout (*Scx-Cre::Bmal1*^*lox/lox*^) mice (n=3 and p=0.6296 for Achilles tendon; n=3 and p=0.7465 for tail tendon). **E**, Relative expression of *Mmp14*, measured at ZT3 and ZT15 in WT and *Clock*Δ19 tail tendons (n=3).

**Figure S2. *Mmp14*-deficient tendons accumulate small-diameter collagen fibrils**. Transmission electron microscopy of control (**A**) and CKO (**B**) mouse tail tendons, at T4 (23 weeks postnatal age). Scale bar, 0.5 µm. Asterisks indicate fibricarriers containing cross-sectional profiles of small-diameter collagen fibrils.

**Figure S3. Depletion or inactivation of *Mmp14* weakens time-dependent differences of protein abundance. A**, Correlation matrix showing Pearson’s R-squared values between datasets resulting from mass spectrometry analysis of tail tendon tissue from CKO and control mice at ZT3 and ZT15 time points within the circadian cycle (n=3 biological repeats; see Fig. 4). Greater similarity was observed between samples at different timepoints than between CKO vs. Ctrl. **B**, Plot showing fold-changes in protein abundance, correlating changes observed in *Mmp14* CKO tendon tissues between ZT15 vs. ZT3 time points, to changes observed in control tissue between corresponding time points. The data showed significant positive correlation, with a Pearson’s R-squared value of 0.2421 (p < 0.0001). However, the difference between the ZT15 vs. ZT3 timepoints was compressed in the *Mmp14* CKO tissues, evidenced by linear fit to the data (shown in red, with gradient lower than y=x), and consistent with a dampened circadian response (n=3 biological repeats).

**Figure S4. Interaction analysis of non-time-dependent matrisome proteins with differential abundance in CKO tendons. A**, Map of reported interactions between some of the matrisome proteins with increased abundance and **B**, decreased abundance in CKO tendons. MMP14 was added to maps to indicate changes that may be directly influenced by *Mmp14* deletion. **C**, Relative abundance of COL1A1 and COL1A2 peptides in control CKO tendons at ZT3 and ZT15.

**Figure S5. Transmission electron microscopy of *Col-r/r* mouse tendon. A**, schematic showing the location of the ¾-¼ MMP cleavage site (arrow) in collagen-I and its absence in the *Col-r/r* mouse. **B**, Electron microscopy images of tendons cross sections of 23 weeks old wild type and *Col-r/r* mice. Scale bars, 0.5 µM. **C**, Fibril diameter distributions of fibrils measured in wild type and *Col-r/r* tendons.

**Figure S6. MMP Inhibition and *Mmp14* weakens circadian entrainment. A**, Dose response curves showing the effects of GM6001 on the viability of Per2::luc iTTFs assessed by Alamar blue. **B**, Representative bioluminescence recordings from DMSO or GM6001 treated Per2::Luc iTTFs, with and without synchronization with dexamethasone. **C**, *Mmp14* mRNA expression detected by Q-PCR 72 hours after transfection of Per2::Luc iTTFs with scrambled (*siScram)* or *Mmp14* targeting siRNA (*siMmp14*), (n=3). **D**, Representative bioluminescence recordings from siScram and *siMmp14* transfected Per2::Luc fibroblasts, after synchronization with dexamethasone. **E**, Sanger sequencing of *Mmp14* in iTTFs derived from the Per2::Luc mouse (control iTTFs) and a single cell clone with *Mmp14* CRISPR-Cas9 knockout. Sequences proximal to the gRNA binding site are shown. **F**, Western blot detection of MT1-MMP in control Per2::luc iTTFs and single cell clones of *Mmp14* CRISPR-Cas9 knockouts.

**Movie 1:** Step-through movie generated from images collected by serial block face-scanning electron microscopy of 23-week-old mouse Achilles tendon. The object in the center of the stack is a cell with long outstretched processes surrounded by collagen fibrils oriented parallel to the long axis of the tendon. Scale bar, 1 µm.

**Movie 2:** Step-through movie generated from images collected by serial block face-scanning electron microscopy of 21-week-old (i.e., 13 weeks post tamoxifen treatment) conditional knockout mouse Achilles tendon. Gross disruption of the tendon matrix has occurred with collagen fibrils misaligned with respect to the tendon long axis and cell processes difficult to identify. Scale bar, 1 µm.

**Table S1** is a list of proteins identified from control and *Mmp14* CKO tendons at ZT3 and ZT15 by mass spectrometry. (XLSX)

**SourceData FS1** contains original blots for Fig. S1.(PDF)

**SourceData F5** contains original gels for Fig. 5. (PDF)

## Data availability

Raw proteomic data was deposited to the ProteomeXchange Consortium via the PRIDE partner repository with the data set identifier PXD031692.

## Experimental procedures

### Mice

The care and use of all mice in this study was carried out in accordance with the UK Home Office regulations, UK Animals (Scientific Procedures) Act of 1986 under the Home Office licence (70/8858) and with German legal guidelines. Ethical approval was granted by the University of Manchester and by the Regierungspräsidium Köln, Germany (NRW authorization 50.203.2-K 37a, 20/05 and AZ2010.A342). The permission included the generation of CKO out animals. All mice were housed in 12-h light/12-h dark cycle (LD) and are indicated where zeitgeber time (ZT) is used, with the exception of wild type C57BL/6 mice housed in 12-h dark/12-h dark cycle (DD) that were used for circadian time (CT) time-series microarray and protein extraction as previously described ^27^. To generate inducible CKO mice in which *Mmp14* is ablated in fibroblasts upon induction with tamoxifen, we crossed mice expressing CreER recombinase under the control of the *Col1a2* promoter (*Col1a2-CreERT2; C57BL/6)* ^*62*^ with C57BL/6 mice carrying the floxed exons (exons 2 to 4) of the *Mmp14* gene ^63^ to produce *Col1a2-CreERT2::Mmp14*^*lox/lox*^ CKO mice. To induce *Mmp14* ablation, mice were fed tamoxifen-formulated chow at 5-weeks old for 4 weeks. To generate mice in which *Bmal1* is ablated in tendon-lineage cells, we crossed mice expressing Cre recombinase under the control of the *Scleraxis* promoter (*Scx-Cre;* C57BL/6) ^64^ with C57BL/6 mice carrying loxP sites flanking the exon encoding the BMAL1 basic helix-loop-helix domain ^65^ bred on a background with *PER2::Luciferase* knock-in ^24^ to produce *Scx-Cre::Bmal1*^*lox/lox*^ KO mice. *Scx-Cre::Bmal1*^*lox/lox*^ KO mice were validated previously ^5^. *Clock*Δ19 mice were as described previously ^66^. *Col1a1-r/r* mice were imported from Jackson Laboratory (*B6;129S4-Col1a1*^*tm1Jae/J*^) ^*36*^. Mice that showed signs of distress or had problems walking were humanely sacrificed using Home Office approved cervical dislocation.

### Tissue culture

Immortalized mouse tail tendon fibroblasts were prepared from Per2::Luc mice as described previously ^5^. To synchronize the internal circadian pacemaker, cells were treated with 100 nM dexamethasone as described previously ^5^. +GM6001 treatment, cell viability

### Fusion protein constructs

Lentivirus expression of *Mmp14 w*as achieved using vectors generated by Vectorbuilder. Murine *Mmp14* coding sequences were expressed under a CMV promoter, with mutation of the *Mmp14* targeting gRNA, AGATCAAGGCCAATGTTCGGAG, PAM site (AGG to AAA). For mCherry tagging the mCherry coding sequence was introduced after amino acid 536 of murine *Mmp14* following the design of Dr. Philippe Chavrier ^67^, these vectors also carried blasticidin resistance gene. Pitx2-GFP expressing cells were cloned into pLenti PGK Neo DEST (Addgene plasmid # 19067 ref^68^), EGFP was cloned at the C terminus of the coding sequence of Pitx2 isoform 1 (Origene; MR227617). Lentivirus particles were generated as previously described ^69^. Stable cells were created by neomycin selection (250 µg/mL) or 5 µg/mL blasticidin.

### CRISPR/Cas9

To generate *Mmp14* knockout fibroblasts, immortalized tail tendon fibroblasts from the Per2::Luc mouse, generated at previously described ^70^, were transduced with lentivirus particles created from vectors co-expressing Cas9 and gRNA (ABMgood 302361140595). *Mmp14* knockout were achieved using vectors encoding the gRNA AGATCAAGGCCAATGTTCGGAG. After puromycin selection (2.5 µg/mL) we isolated single cell clones by FACS and confirmed knockout of *Mmp14* by real-time PCR and western blotting.

### Bioluminescence imaging and recordings

Bioluminescence was recorded as described previously ^5^. Briefly, tendon fibroblasts were cultured on 35 mm culture dish in recording media containing 0.1 mM luciferin (Merck) in the presence of 100 nM dexamethasone (Merck) and real-time quantitative bioluminescence was recorded using lumiCycle apparatus (Actimetrics). Baseline subtraction was carried out using a 24-hr moving average. Amplitude was calculated as peak-trough difference in bioluminescence of the second peak using base-line subtracted data.

### Light microscopy

Live cell super resolution microscopy for was performed using a Zeiss LSM 880 Axio Observer microscope with Airyscan detection system in SR mode using a Plan-Apochromat 63×/1.4 Oil DIC M27 objective (Carl Zeiss, Jena, Germany). Excitation was achieved using the 488⍰nm line from an argon laser and 594⍰nm HeNe laser and emitted light was collected through appropriate filters to eliminate any spill-over. Raw images were Airyscan processed using the default settings and analyzed using ZEN 2.1 software and the image brightness and contrast adjusted using BestFit.

### RNA isolation and Q-PCR analysis

RNA was isolated from Achilles and tail tendons as described previously ^71^. In brief, tissues were washed with ice-cold PBS 3 times and snap frozen in liquid nitrogen with 500 μL TRIzol Reagent (Thermo Fisher Scientific) using a Mikro Dismembrator S (B. Braun Biotech International) at 2000 Hz for 90 seconds two times. After, 500 µL TRIzol was added to the homogenized samples and RNA was isolated according to the manufacturer’s protocol. DNase treatment of RNA was performed using RQ1 RNase-free DNase (Promega) according to manufacturer’s protocol. RNA integrity was assessed by gel electrophoresis and RNA concentration was measured using a NanoDrop 2000 (Thermo Fisher Scientific). TaqMan Reverse Transcription Reagents (Thermo Fisher Scientific) were used for complementary DNA synthesis and SensiFAST SYBR kit reagents (Bioline) were used for Q-PCR. Primers sequences used were: *Col1a1* GCC TGC TTC GTG TAA ACT CC and TTG GTT TTT GGT CAC GTT CA, *Mmp14* AGT GAC AGG CAA GGC TGA TT and GCC CAC CTT AGG GGT GTA AT, *Pitx2* CCT TAC GGA AGC CCG AGT and AAA GCC ATT CTT GCA CAG C, *Plod2* CAA ATC ATA AAA TCG TCT TTG CAG and CCT CTT CAG TGG GTC GAT GT, and *Gapdh* AAC TTT GGC ATT GTG GAA GG and ACA CAT TGG GGG TAG GAA CA. The 2^-ΔΔCt^ method ^72^ was used to analyze relative fold changes in gene expression compared to the first time point. Microarray data for *Mmp14* expression during 48 h was collected previously ^27^ and the data is available on ArrayExpress (accession number E-MTAB-7743).

### Western blotting

Protein purification and western blot analyses were performed as described previously ^5^. Wild type mice were kept in DD for 36 h to allow endogenous circadian rhythm of gene expression. Protein was purified from tail tendons at 4-h intervals. Primary antibodies used were mouse mAb to MT1-MMP (1:500; clone LEM-2/15.8, Millipore), mAb GAPDH (1:10000; clone Gapdh71.1; Merck), and vinculin (1:500; clone hVIN-1, V9131).

### Sample preparation for proteomic analysis

Tail tendons from wild type and *Mmp14* CKO mice were collected at ZT3 and ZT15. Tissues were homogenized with a bullet blender (with 1.6 mm steel beads; Next Advance) at maximum speed at 4 °C for 5 min in 200 μL of SL-DOC buffer (1.1% sodium laurate, 0.3% sodium deoxycholate, 0.5 mM dithiothreitol (DTT) in 25 mM ammonium bicarbonate), supplemented with protease and phosphatase inhibitor cocktails (Roche). Samples were incubated at 4 °C for 5 min, alkylated with 12 μL of 30 mM of iodoacetamide for 20 min at RT, followed by quenching with 12 μL of 30 mM DTT. Samples were centrifuged at maximum speed for 10 min. Supernatants were transferred into LoBind tubes (Eppendorf) and protein concentrations measured using a Millipore Direct Detect spectrometer. A total of 2 μg protein per sample was digested using trypsin beads in accordance with the manufacturer’s protocol (SMART digestion kit, Thermo Scientific). Supernatants were extracted from the beads and 10% formic acid added to adjust to pH 3. Samples were cleaned using an organic phase extraction method: 400 μL ethylacetate (Sigma Aldrich) was added, the resulting solution thoroughly mixed, centrifuged, and the organic phase extracted and discarded; this process was then repeated. Samples were then desalted, in accordance with the manufacturer’s protocol, using POROS R3 beads (Thermo Fisher) and lyophilized in a speed-vac centrifuge (Heto).

### Liquid chromatography-coupled tandem mass spectrometry (LC-MS/MS)

Samples were dissolved in 15 μl of 5% acetonitrile with 0.1% formic acid. Digested samples were analyzed using an UltiMate® 3000 Rapid Separation liquid chromatography system (RSLC, Dionex Corporation) coupled to either Q Exactive HF (Thermo Fisher Scientific) mass spectrometer. Peptides were separated in a gradient of 95% A and 5% B to 7% B at 1 min, 18% B at 58 min, 27% B in 72 min and 60% B at 74 min at 300 nL/min using a 75 mm x 250 μm inner diameter 1.7 μM CSH C18, analytical column (Waters). Peptides were selected for fragmentation automatically by data dependent analysis.

### Proteomics data processing

Spectra from multiple samples were aligned using Progenesis QI (Nonlinear Dynamics) Peak-picking sensitivity was set to 4/5 and all other parameters were left as defaults. Only peptides with charge between +1 to +4, with 2 or more isotopes were taken for further analysis. Filtered peptides were identified using Mascot (Matrix Science UK), by searching against the SwissProt and TREMBL mouse databases. The peptide database was modified to search for alkylated cysteine residues (monoisotopic mass change, 57.021 Da), oxidized methionine (15.995 Da), hydroxylation of asparagine, aspartic acid, proline or lysine (15.995 Da) and phosphorylation of serine, tyrosine, threonine, histidine or aspartate (79.966 Da). A maximum of 2 missed cleavages was allowed. Peptide detection intensities were exported from Progenesis QI as Excel (Microsoft) spread sheets for further processing using code written in-house in MATLAB with the bioinformatics toolbox (R2015a, The MathWorks, USA). Raw proteomic data was deposited to the ProteomeXchange Consortium via the PRIDE partner repository with the data set identifier PXD031692. Fold-change differences in the quantity of proteins detected in different samples were calculated by fitting a linear regression model to take into account inter-sample variation ^73, 74^. Briefly, peptide intensities were logged and normalised by the median intensity; protein fold-changes were calculated using a mixed-effects linear regression model, considering random (peptides, biological replicates) and fixed (controlled variables) effects. Pathway enrichment analysis was performed using the DAVID bioinformatics resource ^75^. Peptide identifications were filtered via Mascot scores so that only those with a Benjamini-Hochberg false discovery rate (FDR) ≥ 0.05 remained. FDR filtering was performed on combined phospho-enriched and non-enriched datasets as described in ^76^. Raw ion intensities from peptides belonging to proteins with fewer than 2 unique peptides (by sequence) per protein in the dataset were excluded from quantification. Remaining intensities were logged and normalized by the median logged peptide intensity. Peptides assigned to different isoforms were grouped into a single “protein group” by gene name. Only peptides observed in at least 2 samples were used in quantification. Missing values were assumed as missing due to low abundance, an assumption others have shown is justified ^77^. Imputation was performed at the peptide level following normalization using a method similar to that employed by Perseus ^78^ whereby missing values were imputed randomly from a normal distribution centered on the apparent limit of detection for this experiment. The limit of detection in this instance was determined by taking the mean of all minimum logged peptide intensities and down-shifting it by 1.6σ, where σ is the standard deviation of minimum logged peptide intensities. The width of this normal distribution was set to 0.3σ as described in ^78^. Fold-change differences in the quantity of proteins detected in different time-points were calculated by fitting a mixed-effects linear regression model for each protein with Huber weighting of residuals (doi: 10.1214/aoms/1177703732) as described in ^77^ using the fitglme Matlab (The MathWorks, USA) function with the formula:

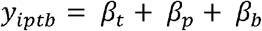

Where y_ipt_ represents the log_2_ (intensity) of peptide p belonging to protein i, at time-point t, in batch b. βs represent effect sizes for the indicated coefficients. Peptide was entered as a random effect whereas time-point was entered as a fixed effect. Standard error estimates were adjusted with Empirical Bayes variance correction according to ^79, 80^. Conditions were compared with Student’s t-tests with Benjamini-Hochberg correction for false positives (doi:10.1111/j.2517-6161.1995.tb02031.x).

### Interactions analysis

Protein interactions were mapped using Pathway Commons version 12 ^81^.

### Collagen content and crosslink analyses

The hydroxylysine aldehyde (pyridinoline) cross-link content of tendon samples was determined by HPLC after acid hydrolysis in 6 N HCl for 24 hrs at 108 °C as previously described ^30^. Results were expressed in residues per mole of collagen based on the hydroxyproline content assayed on the same hydrolysate as a measure of tissue collagen content.

### Electron microscopy

For transmission electron microscopy and morphometric analyses, tendons from mice were prepared as described previously ^29^. Fibril diameter, perimeter and area were measured using ImageJ software to calculate circularity and fibril volume fraction as described previously ^29^. Fibril diameter distributions were plotted and fitted to 3-Gaussian distribution using Sigma Plot 12.0 software (Systat Software) as previously described ^29^. For serial block face-scanning electron microscopy (SBF-SEM), sample preparation and imaging was as described previously using an FEI Quanta microscope fitted with a Gatan 3View® ultramicrotome ^29^. Step-through image stacks were generated using IMOD ^82^ and ImageJ ^83^.

### Recombinant MT1-MMP collagenase assay

Cleanly dissected mouse Achilles tendons were snap frozen in liquid nitrogen, and disrupted in 0.1 M Tris, pH 7.5 using a Mikro-dismembrator S (2 × 90 s, 2000 rpm, cooled with liquid nitrogen) to generate dispersed collagen fibrils. Recombinant human MT1-MMP (250 μl) was dialyzed against four changes of assay buffer (50mM Tris-HCl, 150 mM NaCl, 10mM CaCl_2_, 0.02% NaN_3_, 0.05% Brij35, pH 7.5) at 0°C for 48 h. Pepsin-treated bovine collagen type I (2 µg at 100 μg/ml final concentration; Thermo Fisher Scientific) or a mixture of dispersed collagen fibrils from mouse Achilles tendon in assay buffer (50 mM Tris-HCl, 150 mM NaCl, 10 mM CaCl2, 0.02% NaN3, 0.05% Brij35, pH 7.5) (2 µg at 200 µg/ml final concentration) were incubated with or without rMT1-MMP in assay buffer (20 µl reactions) for 18 h at 30°C and 37°C. Protein concentrations of collagen substrates were confirmed by performing BCA protein assay (ThermoFisher) according to manufacturer’s instructions. Following digestion, samples were separated by 10% SDS-PAGE for mass spectrometry analysis or sampled on a carbon-filmed 400 mesh copper grid for analysis by scanning transmission electron microscopy (STEM).

### Mass spectrometry

Gels stained with Instant Blue stain for 1 h at room temperature. Gels were de-stained in water. Proteins were excised from SDS-PAGE gels and dehydrated using acetonitrile followed by vacuum centrifugation. Dried gel pieces were reduced with 10 mM dithiothreitol and alkylated with 55 mM iodoacetamide. Gel pieces were then washed alternately with 25 mM ammonium bicarbonate followed by acetonitrile and dried by vacuum centrifugation. Samples were digested with trypsin, AspN or GluC overnight at 37°C, as above. Digested samples were analyzed by LC-MS/MS using an UltiMate 3000 Rapid Separation LC (RSLC, Dionex Corporation, Sunnyvale, CA) coupled to an Orbitrap Elite (Thermo Fisher Scientific, Waltham, MA) mass spectrometer. Peptide mixtures were separated using a gradient from 92% A (0.1% FA in water) and 8% B (0.1% FA in acetonitrile) to 33% B, in 44 min at 300 nL/min, using a 250 mm x 75 μm i.d. 1.7 μM BEH C18, analytical column (Waters). Peptides were selected for fragmentation automatically by data dependent analysis.

### STEM and measurement of fibril mass per unit length and calculation of effective hydrated fibril diameter

The mass per unit lengths of the unstained fibrils were measured using annular dark-field scanning transmission electron microscopy (STEM), as described ^35^. STEM images were acquired on an FEI Tecnai 120 TEM (FEI, Eindhoven, Netherlands) fitted with STEM attachments, including a high-angle annular dark-field detector (Fischione, London, UK). A camera length of 350 cm was used to give an angular collection range of 15–75 mrad. Images (1024 × 1024) were acquired at an instrumental magnification of 17,000, corresponding to a pixel size of 5.23 nm, as determined using a diffraction grating replica (2160 lines per mm). Tobacco mosaic virus (a gift from Dr. John Carr, Department of Biochemistry, University of Cambridge, UK) was used as a calibration standard of mass per unit length (131 kDa nm^-1^). The electron dose was kept sufficiently low (2–3 e A^-2^) to produce negligible mass loss. Mass per unit length measurements were measured from STEM images using the Semper6 image analysis software (Synoptics, Cambridge, UK). For type I collagen fibrils the number (Na) of molecules in the transverse section of the overlap zone of each D-period is given by: Na = M_L_/(M/5D) where M_L_ is the mass per unit length of the fibril (kDa / nm), M is the molecular mass (kDa) and D is the axial periodicity (nm). Using values of M=290 kDa for type I collagen and D=67.2 nm (the measured axial periodicity of the unstained, dehydrated fibrils adsorbed on the carbon film) gives: Na = 1.155 x M_L_. Assuming a circular transverse shape for the fibril and a molecular packing density based on X-ray diffraction data from rat-tail tendon ^84^, the effective diameter (W_eff_) of the fully hydrated fibril is then given by: W_eff_ = 1.6 √Na.

### Statistical analysis

Data were evaluated using Two-tailed Student’s t-test, two-way ANOVA or non-parametric, two-tailed Mann-Whitney test. Results were presented as mean ± SEM from at least three independent experiments. Differences were considered significant at the values of *P < 0.05, **P < 0.01 and ***P < 0.001.

