## Supplemental figures 1-6 and source data for "Mmp14 is required for matrisome homeostasis and circadian rhythm in fibroblasts"

#### Figure S1

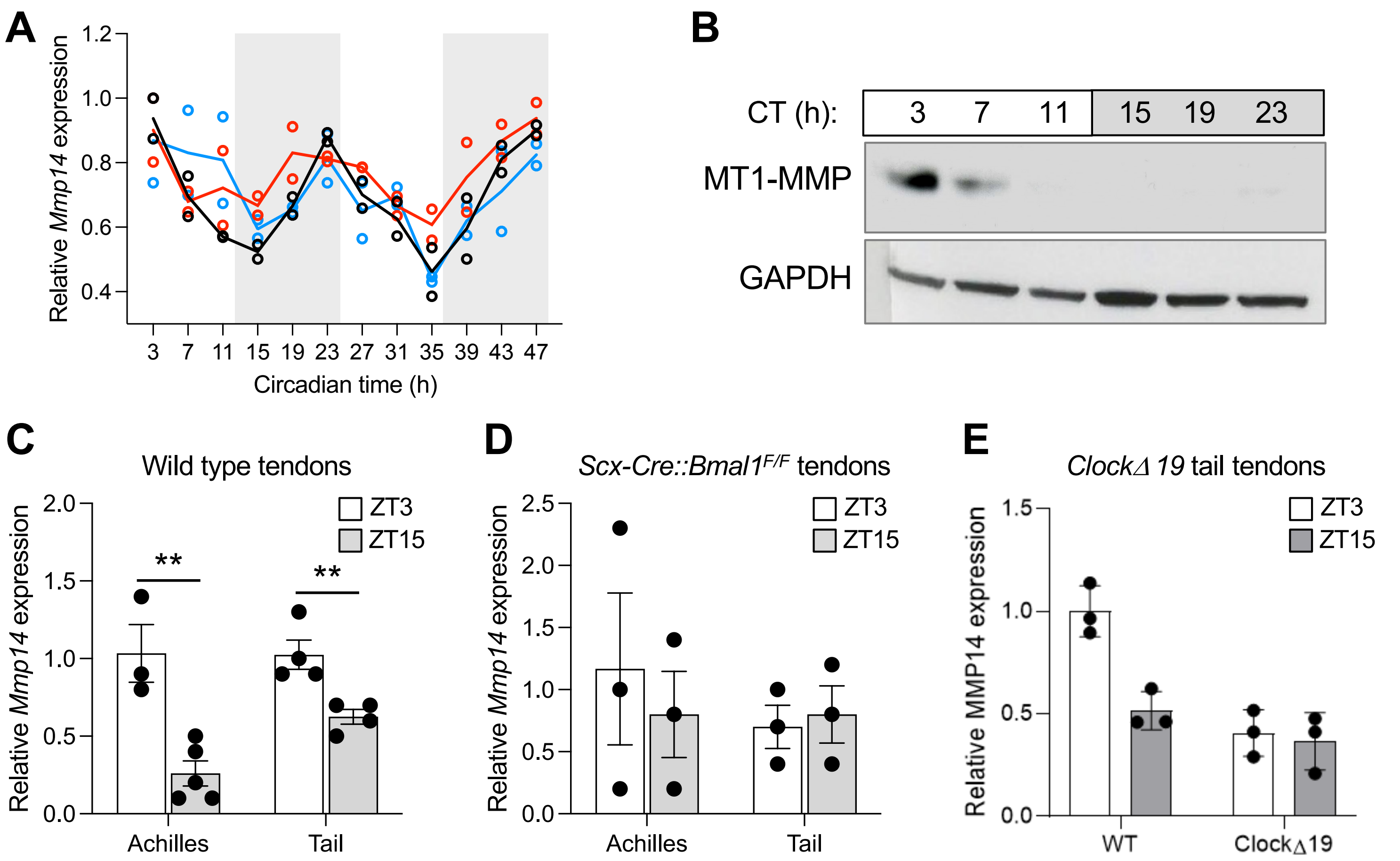


**Figure S1. *Mmp14* expression and MT1-MMP translation in tendon are under circadian clock control. A**, Expression of *Mmp14* mRNA in WT mouse tail tendons analyzed by microarray every 4 h during 48 h, relative to the first time point. Three colored lines indicate the results from 3 probe sets on the microarray chip, n=2; CT, circadian time (free-running time, in constant darkness). **B**, Levels of MT1-MMP in WT mouse tail tendons analyzed by western blotting. Levels of GAPDH protein served as a loading control. **C**, Relative expression of *Mmp14*, measured at zeitgeber time 3 (ZT3; 3 hours into the light phase) and ZT15 in mouse Achilles and tail tendons of WT mice (n=3 and **p=0.0045 for Achilles tendon; n=4 and **p=0.0093 for tail tendon), and **D**, tendon-specific *Bmal1* knockout (*Scx-Cre::Bmal1*$^{lox/lox}$) mice (n=3 and p=0.6296 for Achilles tendon; n=3 and p=0.7465 for tail tendon). **E**, Relative expression of *Mmp14*, measured at ZT3 and ZT15 in WT and *ClockΔ19* tail tendons (n=3).

### Figure S2

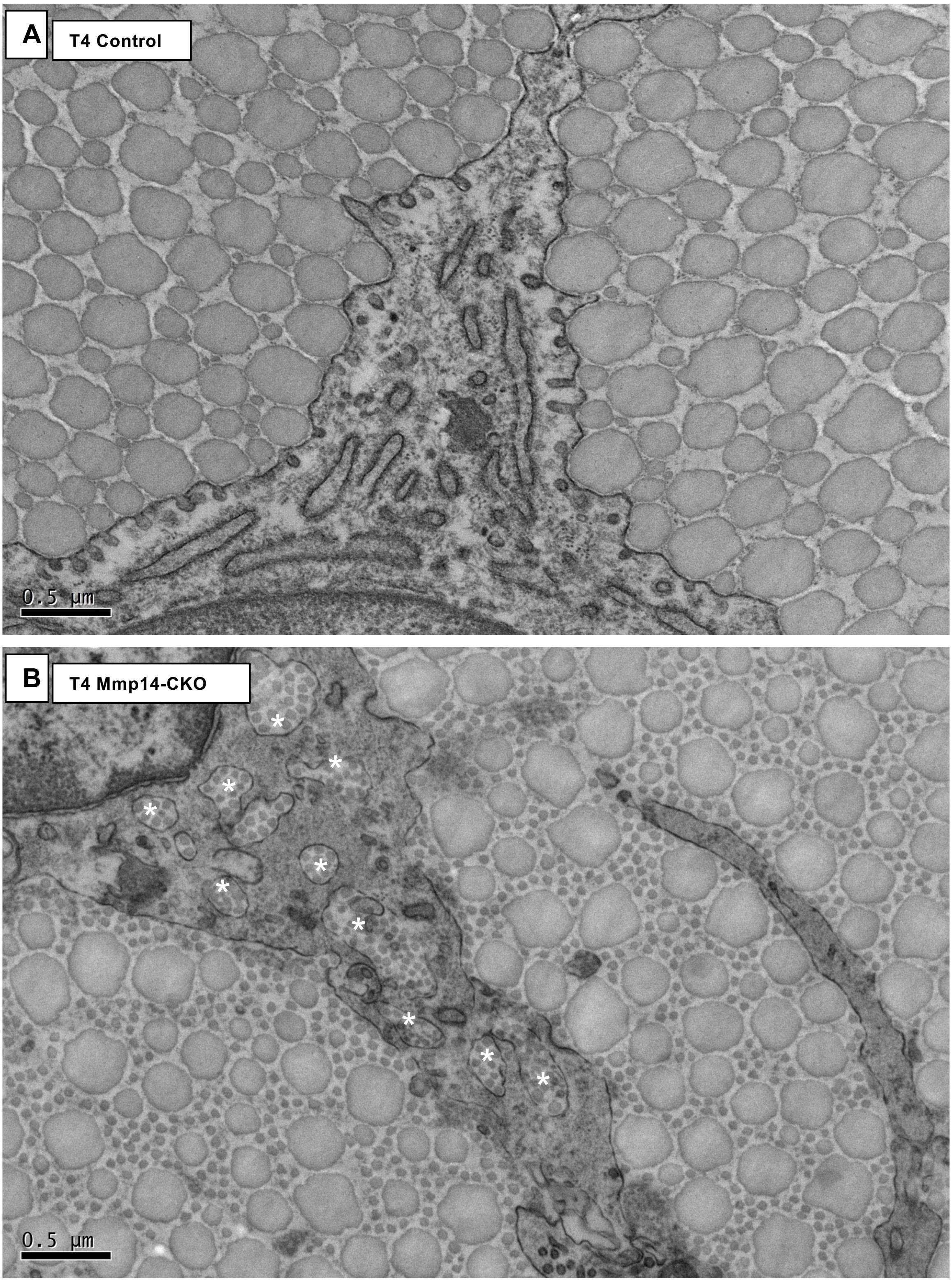


**Figure S2. *Mmp14*-deficient tendons accumulate small-diameter collagen fibrils.** Transmission electron microscopy of control (**A**) and CKO (**B**) mouse tail tendons, at T4 (23 weeks postnatal age). Scale bar, 0.5 µm. Asterisks indicate fibricarriers containing cross-sectional profiles of small-diameter collagen fibrils.

### Figure S3

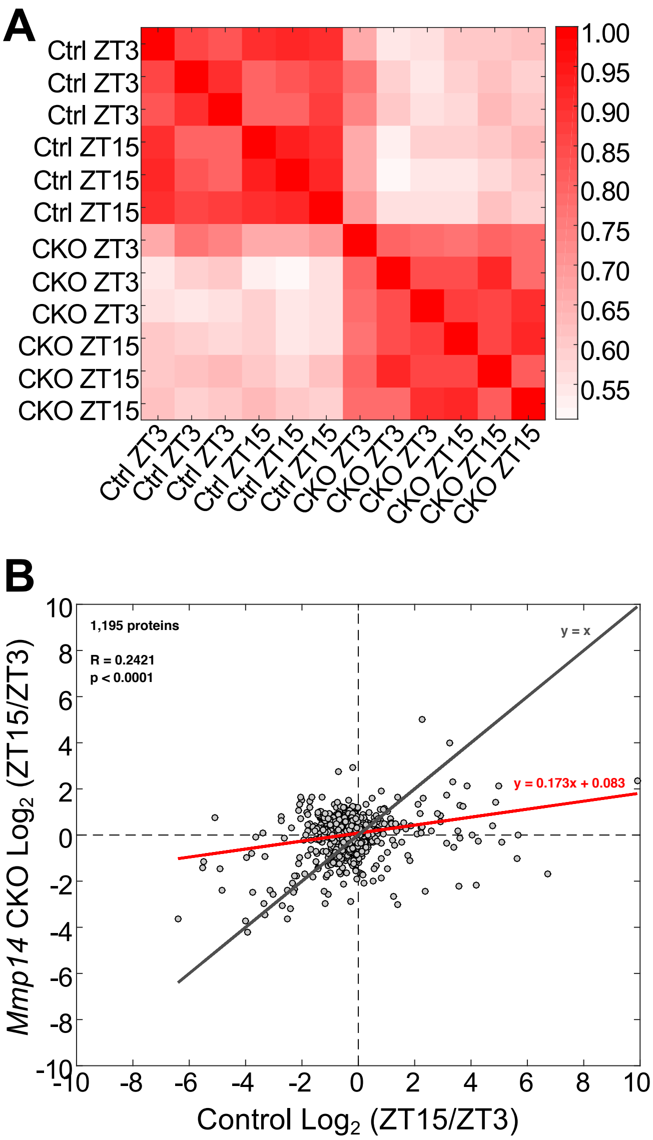


**Figure S3. Depletion or inactivation of *Mmp14* weakens time-dependent differences of protein abundance. A**, Correlation matrix showing Pearson's R-squared values between datasets resulting from mass spectrometry analysis of tail tendon tissue from CKO and control mice at ZT3 and ZT15 timepoints within the circadian cycle (n=3 biological repeats; see Fig. 4). Greater similarity was observed between samples at different timepoints than between CKO vs. Ctrl. **B,** Plot showing fold-changes in protein abundance, correlating changes observed in *Mmp14* CKO tendon tissues between ZT15 vs. ZT3 time points, to changes observed in control tissue between corresponding time points. The data showed significant positive correlation, with a Pearson's R-squared value of 0.2421 ($p < 0.0001$). However, the difference between the ZT15 vs. ZT3 timepoints was compressed in the *Mmp14* cKO tissues, evidenced by linear fit to the data (shown in red, with gradient lower than y=x), and consistent with a dampened circadian response (n=3 biological repeats).

**Figure S4**

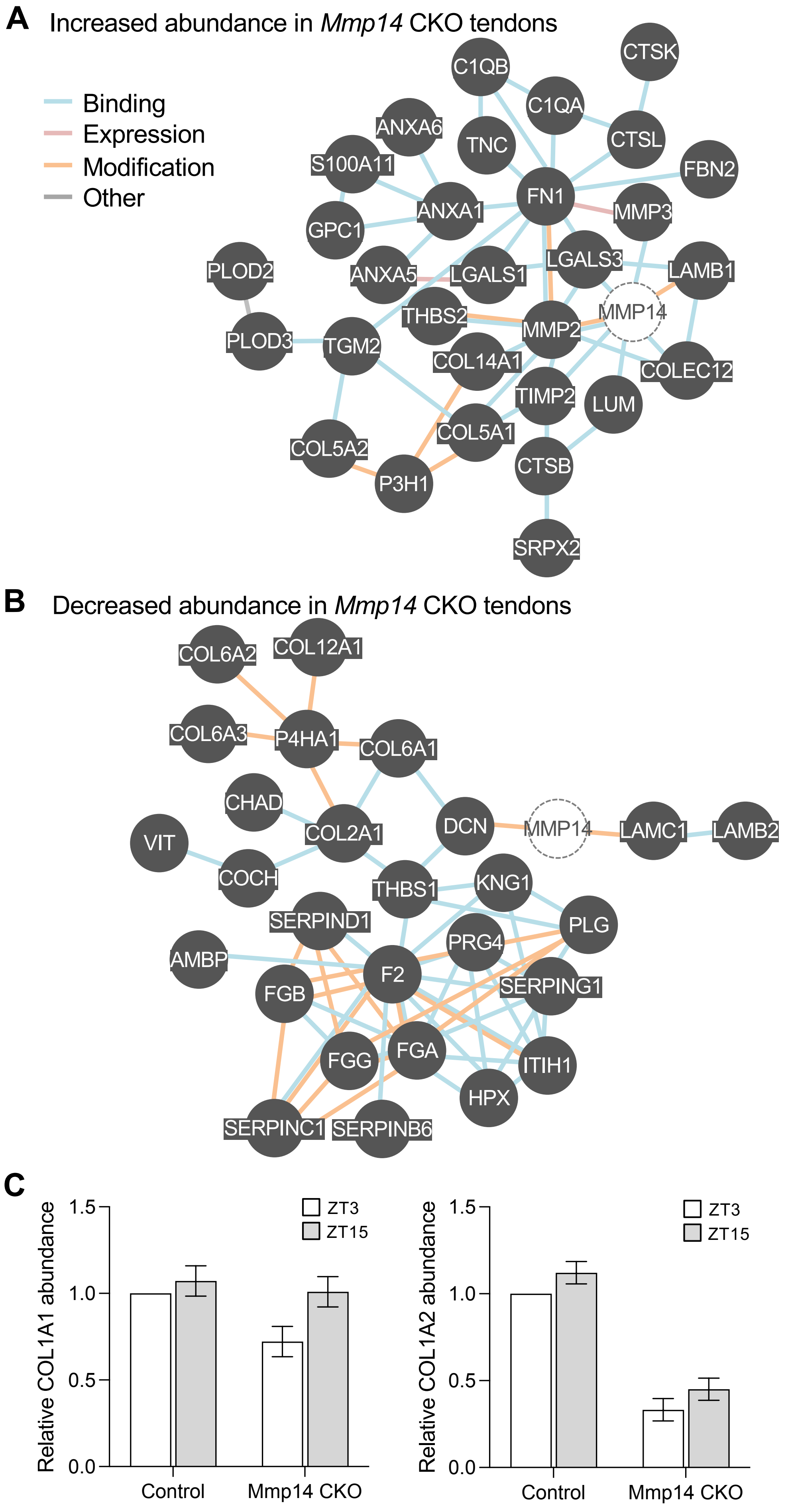


**Figure S4. Interaction analysis of non-time-dependent matrisome proteins with differential abundance in CKO tendons. A**, Map of reported interactions between some of the matrsiome proteins with increased abundance and **B**, decreased abundance in CKO tendons. MMP14 was added to maps to indicate changes that may be directly influenced by *Mmp14* deletion. **C**, Relative abundance of COL1A1 and COL1A2 peptides in control CKO tendons at ZT3 and ZT15.

### Figure S5

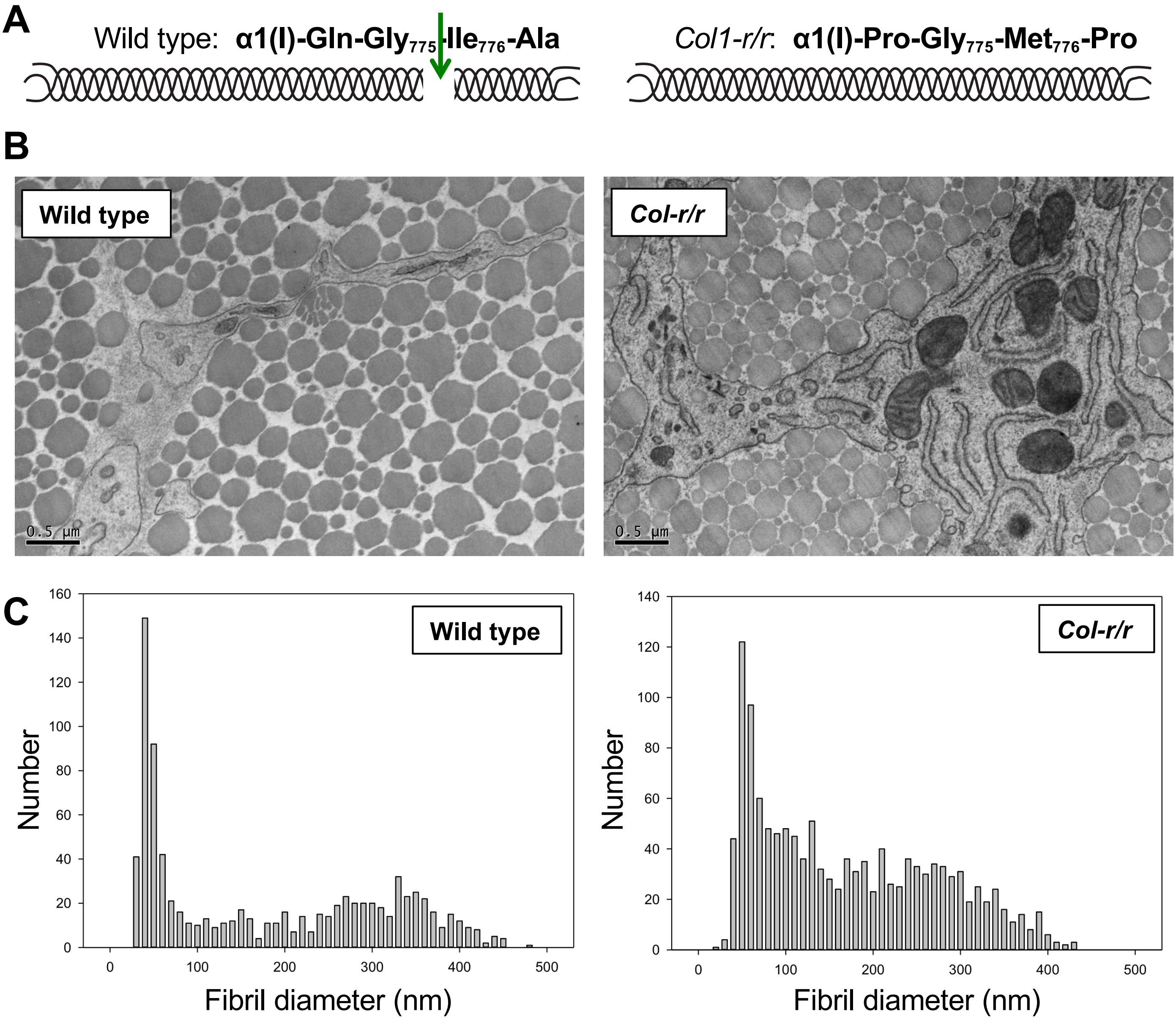


**Figure S5. Transmission electron microscopy of *Col-r/r* mouse tendon. A**, schematic showing the location of the ¾-¼ MMP cleavage site (arrow) in collagen-I and its absence in the *Col-r/r* mouse. **B**, Electron microscopy images of tendons cross sections of 23 weeks old wild type and *Col-r/r* mice. Scale bars, 0.5 µM. **C**, Fibril diameter distributions of fibrils measured in wild type and *Col-r/r* tendons.

**Figure S6**

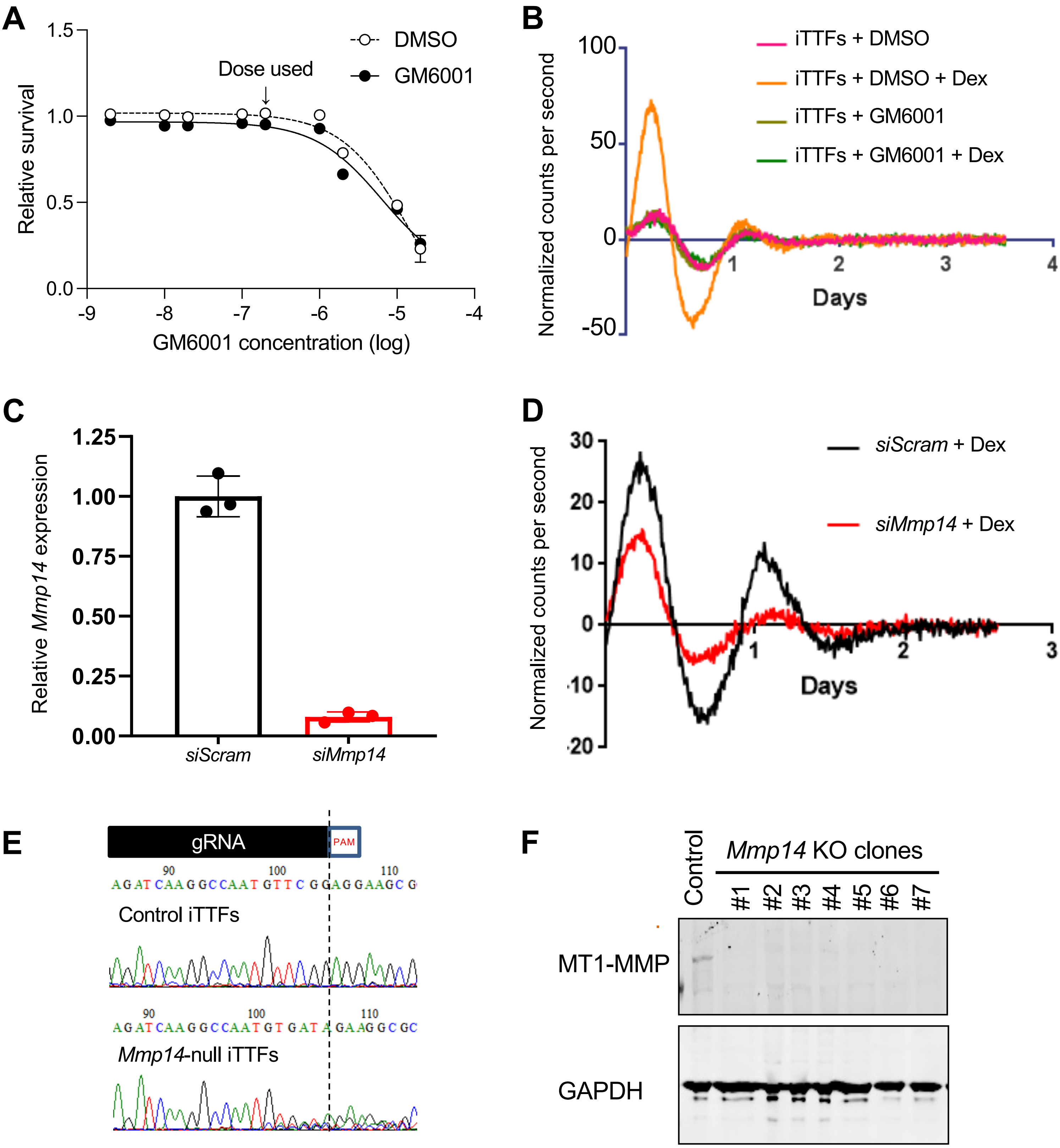


**Figure S6. MMP Inhibition and *Mmp14* weakens circadian entrainment. A**, Dose response curves showing the effects of GM6001 on the viability of Per2::luc iTTFs assessed by Alamar blue. **B,** Representative bioluminescence recordings from DMSO or GM6001 treated Per2::Luc iTTFs, with and without synchronization with dexamethasone. **C**, *Mmp14* mRNA expression detected by Q-PCR 72 hours after transfection of Per2::Luc iTTFs with scrambled (*siScram*) or *Mmp14* targeting siRNA (*siMmp14*), (n=3 biological replicates)). **D**, Representative bioluminescence recordings from *siScram* and *siMmp14* transfected Per2::Luc fibroblasts, after synchronization with dexamethasone. **E**, Sanger sequencing of *Mmp14* in iTTFs derived from the Per2::Luc mouse (control iTTFs) and a single cell clone with *Mmp14* CRISPR-Cas9 knockout. Sequences proximal to the gRNA binding site are shown. **F**, Western blot detection of MT1-MMP in control Per2::luc iTTFs and single cell clones of *Mmp14* CRISPR-Cas9 knockouts.

### Source data for Figure S1

#### Figure S1B

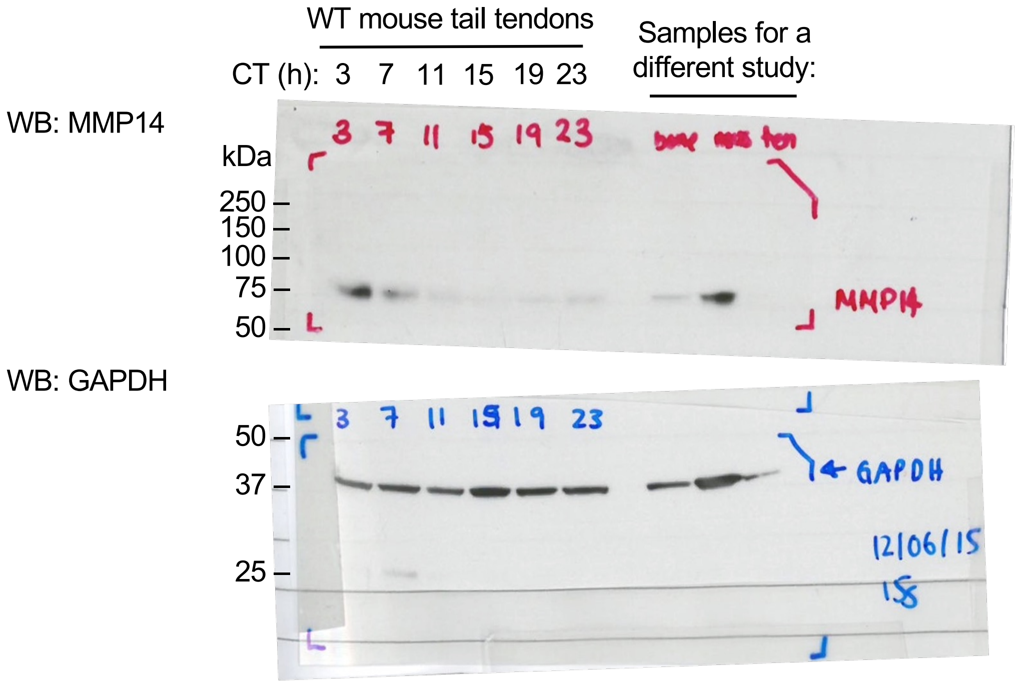

### Source data for Figure 5

#### Figure 5B

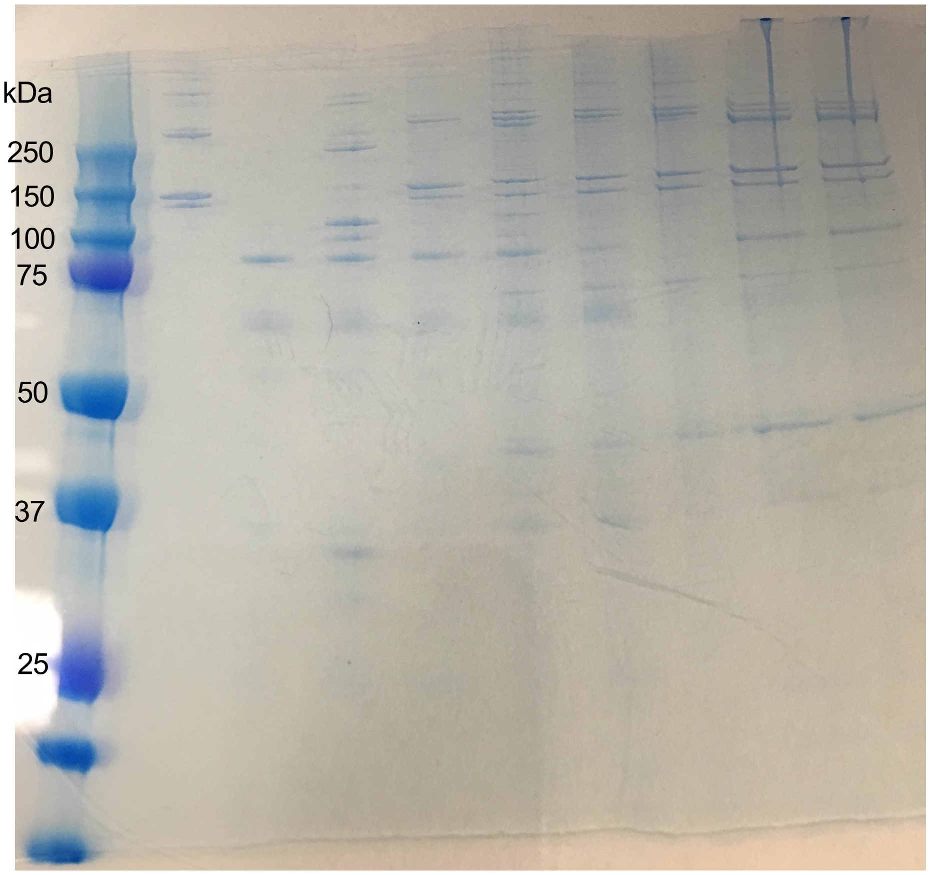
